# The proteolysis of ZP proteins is essential to control cell membrane structure and integrity of developing tubes

**DOI:** 10.1101/2023.07.12.548712

**Authors:** Leonard Drees, Dietmar Riedel, Reinhard Schuh, Matthias Behr

**Affiliations:** Research Group Molecular Organogenesis, Department of Molecular Developmental Biology, Max Planck Institute for Multidisciplinary Sciences, Am Faßberg 11, 37077 Göttingen, Germany; Facility for electron microscopy, Max Planck Institute for Multidisciplinary Sciences, Am Faßberg 11, 37077 Göttingen, Germany; Cell biology, Institute for Biology, Leipzig University, Philipp-Rosenthal-Str. 55, 04103 Leipzig, Germany

**Keywords:** Aneurysms, cell membrane, extracellular matrix, Matriptase, TGFβ, trachea, vessels, Zona Pellucida

## Abstract

Membrane expansion integrates multiple forces to mediate precise tube growth and network formation. Defects lead to deformations, as found in diseases such as polycystic kidney diseases, aortic aneurysms, stenosis, and tortuosity. We identified a mechanism of sensing and responding to the membrane expansion of tracheal tubes. We show in *Drosophila* that Zona Pellucida domain proteins Piopio and Dumpy cooperate to integrate mechanical stress at cell membranes and luminal matrix. When tension appears at the apical membrane due to tracheal tube length expansion, Piopio undergoes ectodomain shedding by the Matriptase homolog Notopleural, which releases Piopio-mediated linkages between membranes and extracellular matrix. Failure of this process leads to deformations of the apical membrane and comprises tubular network function. We also show conserved ectodomain shedding by the human matriptase during TGF-β signaling, both of which are required in the lung, providing novel approaches for in-depth analysis of pulmonary diseases caused by cell and tube shape changes.

## Introduction

Tube networks are essential for organisms transporting liquids, gases, or cells across bodies. Endothelial and epithelial cells generate such networks with strict hierarchical order and precise tube dimensions as a prerequisite for proper tube network functioning (Ochoa-Espinosa and Affolter, 2012; Potente and Mäkinen, 2017). Defective cell shapes and tube dimensions result in severe syndromes such as chronic obstructive pulmonary diseases (384 million people in 2010) with high global mortality as well as tube dysfunctions like aortic aneurysms (152,000 deaths worldwide) in blood vessel systems and polycystic kidney diseases (1 in 1000 people) (Harris and Torres, 2009; Barnes et al., 2015; Adeloye et al., 2015; Quintana and Taylor, 2019; Zhai et al., 2021). One fundamental question regarding the formation of functional tube systems is how cells balance forces at the cell membranes emerging during organ growth and the simultaneous maintenance of the tubular network integrity. *Drosophila melanogaster* embryos form a tracheal system, an excellent model for studying molecular mechanisms controlling cell expansion in combination with tube elongation.

The development of tracheal tubes is subject to precise genetic control. Genes that control cell polarity, junction formation at the lateral membranes, cytoskeletal organization, and intracellular trafficking of tube size determinants support apical cell membrane expansion (Behr et al., 2003; Tonning et al., 2005; Tsarouhas et al., 2007; Laprise et al., 2010; Syed et al., 2012; Förster and Luschnig, 2012; Nelson et al., 2012; Dong et al., 2014; Olivares-Castineira and Llimargas, 2017; McSharry and Beitel, 2019; Skouloudaki et al., 2019). In contrast, genes controlling the meshwork of chitinous apical extracellular matrix (aECM) formation restrict excessive cell expansion to prevent tube overexpansion (Moussian et al., 2006; Wang et al., 2006; Luschnig et al., 2006; Petkau et al., 2012; Tiklova et al., 2013; Öztürk-Çolak et al., 2016). Subsequently, genetically controlled mechanisms establish tracheal airway clearance, aeration, and tube stabilization (Tsarouhas et al., 2007; Behr et al., 2007; Stümpges and Behr, 2011; Drees et al., 2019; Tsarouhas et al., 2019; Behr and Riedel, 2020).

Previous studies revealed that axial and radial forces affect tracheal tube elongation. The apical membrane grows axially, pulling on the associated aECM until the aECM’s elastic resistance balances the elongation force throughout the tubes (Dong and Hayashi, 2015). Given the association between cell membranes and aECM, ongoing tube expansion and luminal shear stresses inevitably lead to problematic membrane tension. Once forces are out of equilibrium, tracheal tubes show curvy appearance. Similarly, also blood vessels can appear unstable, twist, kink, and buckle. High stresses even lead to vascular damage and aneurysm rupture (Han et al., 2013; Dong et al., 2014). However, it is not known how cells manage to integrate the axial forces to stabilize the cell membrane and aECM.

Zona pellucida (ZP)-domain proteins are critical components of apical cell membranes and aECM (Plaza et al., 2010) and assemble into extracellular fibrillar polymers (Jovine et al., 2005; Litscher and Wassarman, 2020). For example, Uromodulin plays a role in chronic kidney diseases and hypertension (Rampoldi et al., 2011). Secreted Uromodulin requires proteolysis at the apical cell membrane for shedding and polymerization within the tube lumen (Brunati et al., 2015). Similarly, the ZP domain protein Piopio (Pio) (Fig. S1A) is secreted into tracheal tubes of *Drosophila* embryos (Grieder et al., 2008; Massarwa et al., 2009). Pio restricts the elongation of autocellular junctions (Jaźwińska et al., 2003) and anchors Spastin, a microtubule-organizing protein, to the apical membrane (Brodu et al., 2010). However, while Pio is a transmembrane protein, it was detected in the tube lumen (Jaźwińska et al., 2003), but the mechanism of its release remains unknown. A promising candidate for Pio proteolysis is Notopleural (Np), the functional homolog of the human Matriptase. It is a type-II single-transmembrane serine protease in tracheal and lung epithelia and is capable of ectodomain shedding (Bugge et al., 2009). Initial *in vitro* studies prove that Np cleaves the Pio ZP domain (Drees et al., 2019). The elastic luminal matrix is essential for the integrity of the tubular network. The matrix balances elongation forces in the anterior-posterior direction during tube elongation (Dong et al., 2014). Here we show that the ZP domain proteins Pio and, Dumpy, as well as the protease Np sense and support balancing of mechanical stresses when tracheal tubes elongate, to ensure normal membrane-aECM morphology.

## Results

### Pio maintains structural cell membrane continuity

The tracheal lumen matrix consists of a viscoelastic material that is coupled to the apical membrane. The precise balance between apical membrane growth and luminal matrix resistance determines tube shape (Dong et al., 2014). Based on these observations, we expect the following scenario: first, apical membrane growth and opposing restriction by the extracellular matrix produce increased tensile stress during tube expansion of stage 16 embryos (Fig. 1A). Second, increasing tension impacts membrane-matrix couplings, which provide the proper balance between both. Thus, enhanced tensile stress may lead to either release or remodeling of the membrane-matrix couplings to avoid potential deformations. The Pio protein contains a transmembrane and an extracellular ZP domain, suggesting that it may link tracheal cells to the aECM. Using CRISPR/Cas9, we generated three *pio* lack of function alleles (Fig. S1A), all exhibiting embryonic lethality and identical tracheal mutant phenotypes. In the following analyses, we focused on *pio^17c^*. The *pio* null mutant embryos revealed the dorsal and ventral branch disintegration phenotype previously described using a *pio* point mutation (Jaźwińska et al., 2003). However, the *pio* null mutant embryos of stage 16 showed over-elongated tracheal dorsal trunk tubes (see below). We compared the dorsal trunk morphology between control and *pio* mutant embryos by using the septate junction (SJs) marker Megatrachea (Mega) (Behr et al., 2003). The early stage 17 control embryos revealed tight appearance of tracheal cells and adjacent luminal extracellular matrix. In contrast, the corresponding *pio* mutant embryos showed irregular bulge-like gaps between the Mega-marked cell membrane and apical matrix (Fig. 1B). Such gaps were not detectable in late-stage 15 wild-type (*wt*) or *pio* mutant embryos (Fig. S1B). This suggests that the gaps arise in stage 16 *pio* mutant embryos during tube length expansion.

**Figure 1:**
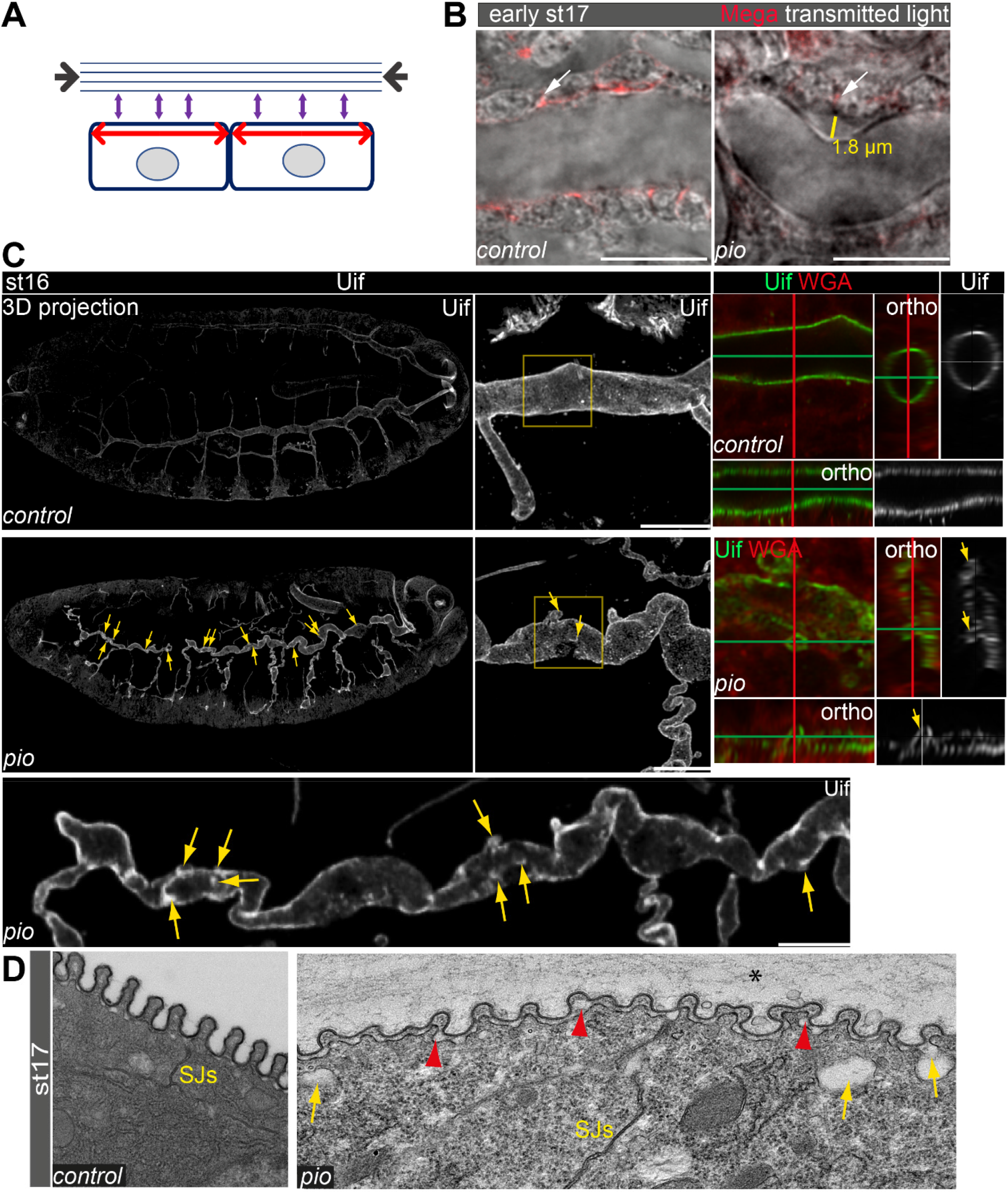
Pio supports structural continuity of the apical cell membrane. **(A)** Model implicates the axial and longitudinal forces (arrows) acting apical on cell membrane and extracellular matrix of stage 16 embryos when tracheal lumen expands in growing tubes. **(B)** Confocal images of *wt* and *pio* mutant stage (st) 17 embryos. Lateral membrane is marked by Megatrachea (Mega; red, arrows) immunostaining, and transmission light visualizes the tracheal cells and lumen. The yellow line indicates the distance between the apical cell surface and the detached luminal aECM. **(C)** Confocal LSM Z-stack (overview) and Airyscan (close up) images of immunostainings are displayed as maximum intensity (3D projection) and orthogonal (ortho) projections using Uif antibody and WGA. Scale bars indicate 10µm. Stage 16 control embryo showed straight apical cell membrane and tracheal tubes. All *pio* null mutant (n=10) embryos revealed curly elongated tracheal tubes and unusual bulges of the apical cell membrane (yellow arrows in overview and close up). Left images show focus on the region (marked by the yellow frame in close up) of orthogonal analyses of the dorsal trunk and corresponding projections are displayed in middle and right images. Note the membrane bulges interfere with the tube lumen integrity (ortho-target cross in mutant). Control and *pio* mutant embryos were fixed and stained together. **(D)** TEM analysis of late-stage 17 *wt* embryos reveal SJs and chitin-rich taenidial folds. The corresponding *pio* mutant embryos (n=4) showed normal SJs formation but unusual apical cell membrane deformations (yellow arrow), reduced chitin (red arrowheads), taenidial folds with disorganized pattern, and extracellular matrix material within the tube lumen (*).

To study the role of Pio during tube length expansion, we examined *pio* mutant stage 16 embryos using the apical cell membrane marker Uif. This revealed unusual apical cell membrane deformations most prominent at the dorsal trunk (Fig. 1C). Corresponding control embryos of the same fixations and stainings did not show any tube size defects and membrane bulges (Fig. 1C). Additional analysis of orthogonal projections along confocal axes revealed straight apical membranes of tracheal tubes in the control embryos while comparable *pio* mutants contained numerous small membrane deformations. Moreover, these membrane deformations compromised normal tube lumen shape (Fig. 1C; orthogonal projections). Furthermore, the ultrastructural analysis of stage 17 *pio* mutant embryos confirmed the apical cell membrane deformation with unusual gaps between the membrane and aECM, while control embryos lacked such gaps (Fig. 1D; Fig. S1C). These results indicate that Pio is required to stabilize or maintain structural membrane-matrix formation.

Anti-Pio antibody (kindly provided by Markus Affolter), detects a short stretch within the Pio ZP domain (Jaźwińska et al., 2003)). Immunostainings confirmed Pio protein expression in the tracheal system of stage 16 embryos when tubes expand (Fig. 2A). In these embryos, tracheal Pio staining is detectable at the membrane and is enriched within the tracheal lumen, consistent with previous findings (Massarwa et al., 2009; Dong et al., 2014). At the membrane of st16 control embryos, Pio overlaps at discrete points with the apical cell membrane determinant Crumbs (Crb) and wheat germ agglutinin (WGA) that visualizes the cell membranes (Fig. S2A,B). In *crb* mutant embryos, the apical membrane is compromised (Laprise et al., 2010), but luminal content is still secreted (Stümpges and Behr, 2011; Olivares-Castiñeira and Llimargas, 2018). These *crb* mutant embryos lacked Pio staining at the membrane, and instead, Pio concentrated within the tube lumen. Control embryos showed normal Pio distribution at the membrane (Fig. 2B).

**Figure 2:**
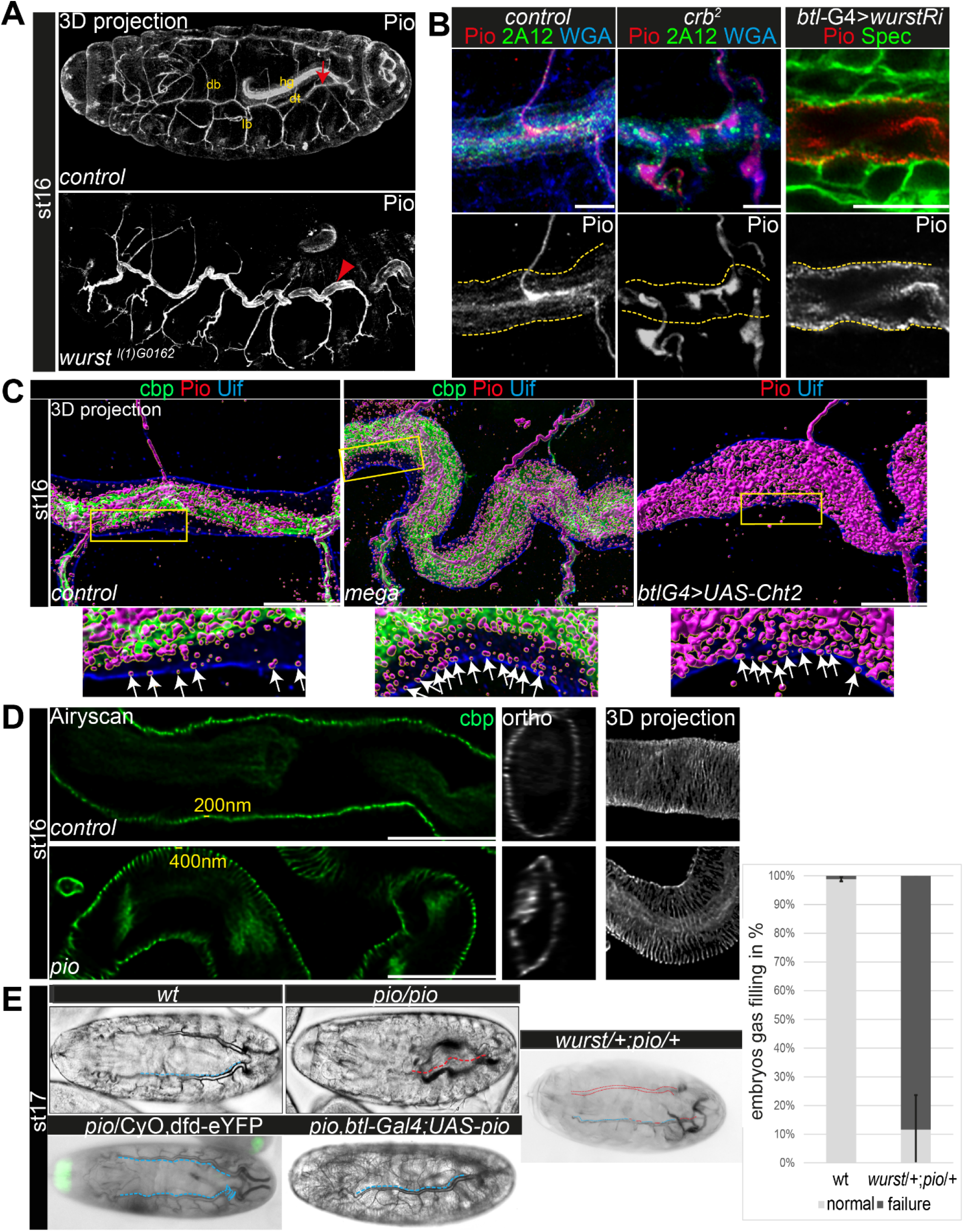
Pio localization depends on the apical membrane and supports tracheal air-filling. Maximum intensity projections of confocal Z-stacks are shown in A, single confocal images in B, and Airyscan images in C,D, respectively. Bright filed microscopy is shown in E. Scale bars indicate 10µm. **(A)** Pio protein is expressed in tracheal tubes (db, dorsal branches; lb, lateral branches; dt, dorsal trunk) and other ectodermal epithelial organs. Pio accumulates in the tracheal lumen (arrow) of control embryos. In contrast, Pio staining showed unusual accumulation at the apical cell membrane of *wurst* mutant embryos (arrowhead point to Pio accumulation). It is of note that embryos were stained together. Shown are whole-mount embryos. hg, hindgut **(B)** In *crb* mutant embryos, Pio staining is within the luminal matrix but not at the apical cell membrane (indicated by yellow dashes). Corresponding control embryos showed Pio at the membrane and predominantly within the lumen. The tracheal-specific *wurst* knockdown shows Pio accumulation at the apical cell membrane similar to the *wurst* mutant embryo (compare with A). 2A12 detects the aECM protein Gasp. WGA labels cell membrane surfaces and chitin predominantly. **(C)** Imaris 3D projection. In wt Pio (magenta) is detectable in a punctuate pattern at the tracheal apical cell membrane, partially overlapping with Uif (blue) in stage 16 control embryos. Chitin- binding probe (cbp; green) labels chitin. The stage 16 *mega* mutant embryo (n=5) showed Pio accumulation at the apical cell surface. The tracheal expression of the Chitinase 2 (n=5) showed a disturbed Pio pattern, including accumulation of Pio puncta at the apical cell surface. The lower panel shows close-ups of the apical cell membrane of the framed area in the upper images. The white arrows indicate Pio puncta at the Uif marked apical cell membrane. We chose comparable regions where a gap formed between the cell membrane and chitin matrix to detect the apical Pio puncta. Note that *mega* mutant embryo (20 Pio puncta) contains twice as much Pio puncta at the Uif stained apical cell membrane as the control (9 Pio puncta); both show tracheal metamers 7-9. **(D)** Airyscan images (left) and orthogonal (middle) and 3D projections (right) of control and *pio* mutant late-stage 16 embryos. cbp detects chitin at the taenidial folds and within the tracheal lumen of control and *pio* mutant embryos. The *pio* mutant embryos show loose taenidial fold patterns with enlarged distances between the ridges. **(E)** Late-stage 17 *wt* embryos revealed normal tracheal air-filling (indicated with blue dashes). All *pio* (n=20) mutants embryos and 89% of transheterozygous *wurst;pio* (n=10) mutant embryos showed tracheal air-filling defects (red dashes indicate liquid filled airways), while almost none of the control embryos showed defects (n=100). Tracheal air-filling defects in embryos were displayed as dark grey bars, and normal air-filling as light grey bars. Error bars indicate the standard deviation. Heterozygous *pio* mutant embryos (n=10) as well as the *btl-Gal4* driven tracheal expression of Pio in *pio* mutant embryos (n=10) revealed normal tracheal air-filling at the end of stage 17 (indicated with blue dashes). Green signal indicates eYFP expression from the balancer chromosome in the heterozygous *pio* mutant embryo.

These observations prompted us to address whether Pio misdistribution depends on apical cell membrane organization. Crb genetically interacts with *wurst* on late airway maturation, including gas-filling (Stümpges and Behr, 2011). The transmembrane protein Wurst (DNJAC22) is a critical component of clathrin-mediated endocytosis (Behr et al., 2007) and controls the internalization of proteins at the apical membrane (Stümpges and Behr, 2011). Stage 16 *wurst* mutant embryos and tracheal-specific wurst RNAi (interference) knockdown embryos revealed unusually increased Pio accumulation at the apical cell membrane compared to control embryos (Fig. 2A; Fig S2B,C).

These findings suggest Wurst-mediated internalization of Pio and raise the question of how intracellular Pio trafficking may occur. Retromer and ESCRT-mediated endosomal sorting regulate key proteins of tracheal tube size control (Dong et al., 2014). Pio is not affected by the retrograde transport from endosomes to the trans-Golgi network (Dong et al., 2013). Vps32/Shrub is a subunit of ESCRTIII, which regulates endocytic sorting of membrane-associated proteins leading to lysosomal degradation and is known to be involved in tracheal tube size control in stage 16 embryos (Dong et al., 2014). Loss of *shrub* leads to the formation of swollen endosomes that accumulate Crb within tracheal cells (Dong et al., 2014). We observed intracellular Crb staining that overlapped with Pio in *shrub* mutant embryos (Fig. S2A). Additionally, Crb and Pio overlapped at the apical cell membrane, suggesting that newly synthesized Pio was secreted as it is known for Crb (Dong et al., 2014). These results indicate that Pio localization relies on apical membrane formation, turnover, and intracellular protein trafficking.

To understand the distribution of Pio under different stress situations, we examined mutants that either increase the apical cell surface or cause severe chitin defects. We generated voxel data of 3D Pio staining projections (Imaris, 3D surface rendering) using Airyscan Z-stacks and deconvolution (SVI Huygens). These data identified Pio voxel in punctuate pattern at and next to the apical cell membrane in *wt* embryos. However, most voxel accumulated around the inner luminal chitin matrix structure (Fig. 2C), resembling the confocal pattern of control embryos (Fig. 2A,B). The disruption of SJs in *mega* mutant embryos caused apical cell surface expansion and increased tube length in stage 16 embryos (Behr et al., 2003). These *mega* mutant embryos showed Pio localization at the chitin matrix but increased staining at and near the apical cell membrane (Fig. 2C; Fig. S3). The ectopic expression of the *chitinase 2 (cht2)* in tracheal cells leads to excessive tube dilation and length expansion due to strongly reduced luminal chitin (Behr et al., 2003; Tonning et al., 2005; Petkau et al., 2012). Also, the tracheal *cht2* expression led to increased Pio staining at and near the apical cell membrane (Fig. 2C; Fig. S3), suggesting that Pio localization is affected by the membrane morphology and the chitin matrix formation.

Next, we investigated if Pio affects the tracheal airway function. First, orthogonal projections of confocal Z-stacks of *pio* mutant embryos revealed that membrane deformation compromised the normal tube lumen (Fig. 1C). Second, the ultrastructure analysis of stage 17 embryos revealed reduced chitin and deformed taenidial folds in *pio* mutant embryos while control embryos established chitin-loaded taenidial folds at the apical cell surface (Fig. 1D; Fig. S1C). It is of note that also confocal microscopy revealed reduced tracheal chitin staining in stage 16 *pio* null mutant embryos (see below). Third, in contrast to *wt*, *pio* mutant st17 embryos showed an irregular pattern of taenidial folds in ultrastructure and Airyscan analysis (Figs. 1D, 2D; Fig S1C). Fourth, the ultrastructure images revealed aECM remnants in the airway lumen of *pio* mutant stage 17 embryos, while control embryos cleared their airways (Fig. S1C). Consistently, the *in vivo* analysis of airways in stage 17 *pio* mutant embryos revealed lack of tracheal air-filling (Fig 2E). The pan-tracheal expression of Pio in *pio* mutant embryos rescued the lack of gas filling (Fig. 2E). Thus, *pio* mutant embryos showed impaired tube lumen clearance of aECM, which prevented subsequent airway gas-filling.

Since Pio localization depends on Wurst, we addressed putative genetic interaction by investigating tracheal air-filling in st17 transheterozygous *wurst* and *pio* mutant embryos. Heterozygous *wurst* and *pio* mutant control embryos showed normal tracheal air-filling (Fig 2E) (Behr et al., 2007). In contrast, transheterozygous embryos bearing one copy of the *pio* and one of the *wurst* mutant alleles showed air-filling defects (Fig. 2E), suggesting genetic interaction. However, *wurst* mutant embryos failed to clear luminal matrix content (Behr et al., 2007), including Pio, thus preventing airway clearance at stage 17 (Fig. S2D). In contrast, the *pio* mutant embryos showed tracheal lumen clearance defects of aECM in ultrastructure analysis (Fig. 1D), but some luminal material, such as the aECM proteins Obstructor-A (Obst-A) and Knickkopf (Knk), are removed from the lumen (Fig. S4A). These findings indicate that Pio function compromises tube airway function.

The chitin-binding protein Obst-A, chitin deacetylases Serpentine (Serp) and Vermiform (Verm), as well as chitin-protein Knk restrict tube expansion (Wang et al., 2006; Moussian et al., 2006; Luschnig et al., 2006; Petkau et al., 2012). In the *pio* mutant embryos, luminal Serp and Verm staining appeared reduced but showed *wt*-like distribution (Fig. S5). Thus, Pio does not control tracheal chitin-matrix secretion, formation, and organization but may affect their maintenance in stage 16 embryos.

Apical cell membrane growth is another essential cellular mechanism of tube growth (Laprise et al., 2010; Dong et al., 2014). However, the apical cell membrane marker Uninflatable (Uif) (Zhang and Ward, 4th., 2009) showed *wt*-like localization in *pio* mutant trachea (Fig. 1C). Further, Crb immunostainings showed normal localization in *pio* mutant embryos (Fig. 3A). Also, the confocal Megatrachea immunostainings and appearance of SJs in ultrastructure analysis were similar between control and *pio* mutant embryos (Fig. 1B,D; Fig. S1B,C). Finally, adherens junctions (AJs) revealed a *wt*-like appearance in *pio* mutant embryos in ultrastructure images (Fig S1C) and also the Armadillo (Arm; *Drosophila* β-Catenin) immunostainings showed *wt*-like pattern at the apicolateral membrane of tracheal cells (Fig 3B). These data show that Pio is not involved in either apical polarity nor in lateral membrane formation, which is consistent with previous findings (Brodu et al., 2010). However, we observed an increased distance in the axial direction between the AJs of the dorsal trunk fusion cells. In addition, we determined an enlarged apical cell surface area due to unusual cell elongation in the axial direction (Fig. 3B-D). These findings indicate that Pio is required to prevent excessive apical cell growth and membrane deformation.

**Figure 3:**
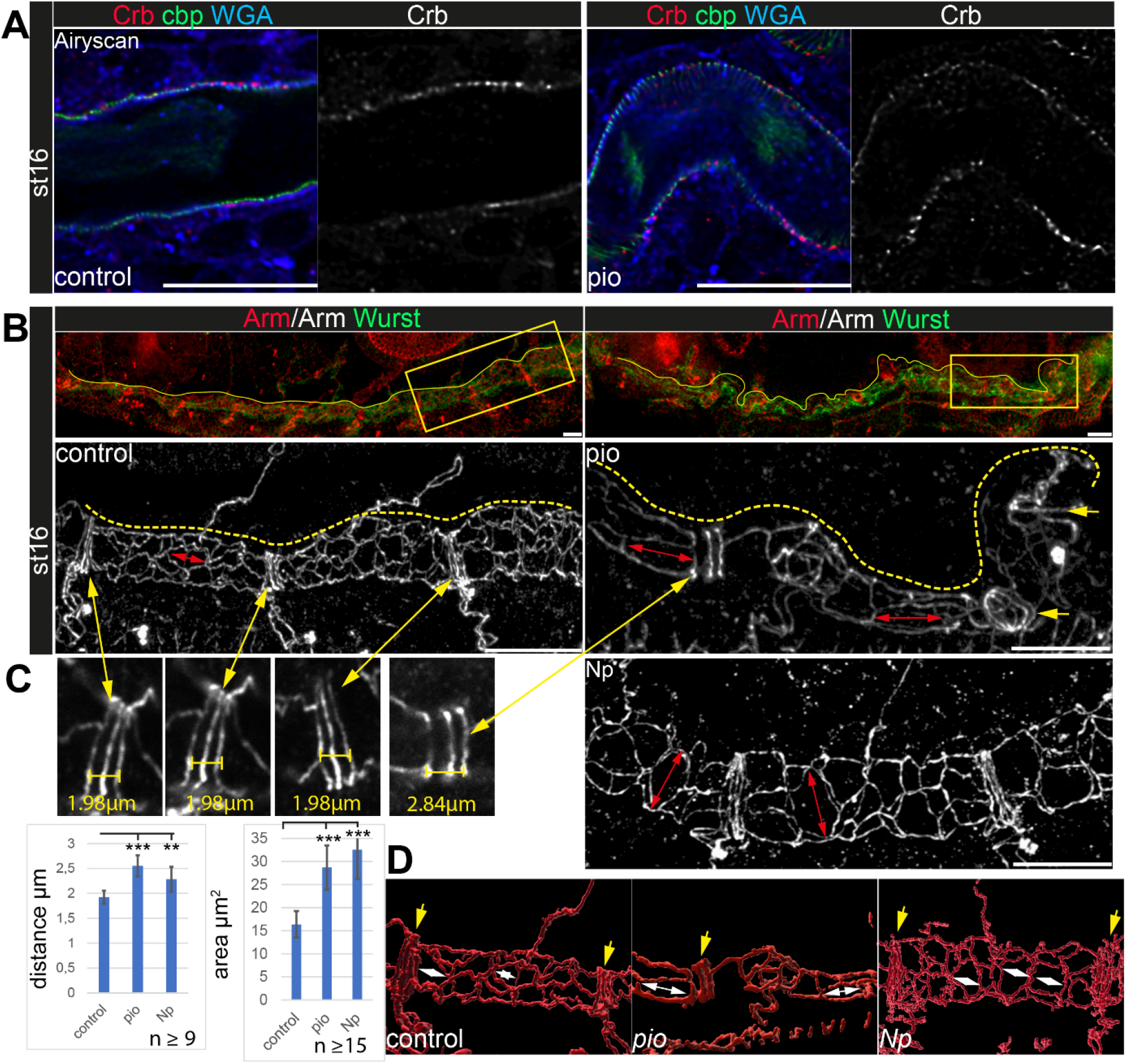
Apical polarity and AJs localization in *pio* and *Np* mutant embryos. Confocal LSM Z-stacks of tracheal dorsal trunk show single layer (A) and 3D projections (B,C) of stage 16 and stage 17 embryos. **(A)** Control and *pio* mutant late stage 16 embryos show Crb (red) staining at the apical membrane, the cell surface marker WGA (blue), and cbp (green) in the tracheal aECM at the apical cell surface and in the luminal cable-like ECM. A single Crb channel is indicated in grey. **(B)** Maximum intensity projections of confocal Z-Stacks of control and *pio* mutant late stage 16 embryos and *Np* mutant early stage 17 embryo. Upper panels show immunostainings with Armadillo (red) and Wurst (green) at the dorsal tracheal trunk. Yellow dashes mark the tracheal tube. Magnifications of the framed regions in the top panel show Armadillo staining in grey (bottom). Red double arrows indicate tracheal cells. Yellow arrows point to AJs of fusion cells **(C)** Yellow double arrows point to magnifications of the Armadillo staining of dorsal trunk fusion cells. The distance of AJs of fusion cells in control, *pio,* and *Np* mutant embryos are indicated in representative images. Plots show AJ distances of fusion cells (n>9) in µm and apical cell area in µm^2^ (n>15). Bars represent ±SD and p-values for AJs distance (*pio* p=5.8e-5; *Np* p=0.0022) and cell area (*pio* p=1.6e-6, *Np* p=2,5e-10), unpaired t-test. **(D)** 3D reconstruction (Imaris surface rendering) of confocal Armadillo immunostainings marking the AJs of control, *pio,* and *Np* mutant embryos. Yellow arrows point to AJs of fusion cells; white double arrows indicate cell length in the axial direction.

To understand Pio dynamics in living embryos, we generated CRISPR/Cas9 mediated homology- directed repair *pio^mCherry::pio^* embryos (Fig S4B). The mCherry::Pio expression in stage 16 embryos (Fig. S4C) resembled Pio protein expression pattern (Fig. 2A). The early-stage 16 embryos showed mCherry::Pio assembling at the tracheal apical cell membranes (Fig. 4A). When tube expansions stopped at the end of stage 16, tracheal mCherry::Pio signal shifted towards the lumen which is distinct of the chitin matrix pattern (Fig. 4A; Video S1). FRAP experiments demonstrated fast recovery of mCherry::Pio expression after photobleaching (Fig. 4B; Video S2). In stage 16 *wt*- like control embryos, mCherry::Pio reappears some minutes after bleaching within tracheal cells and tube lumen, demonstrating that dynamic mCherry::Pio relocation in tracheal cells in the lumen during tube expansion.

**Figure 4:**
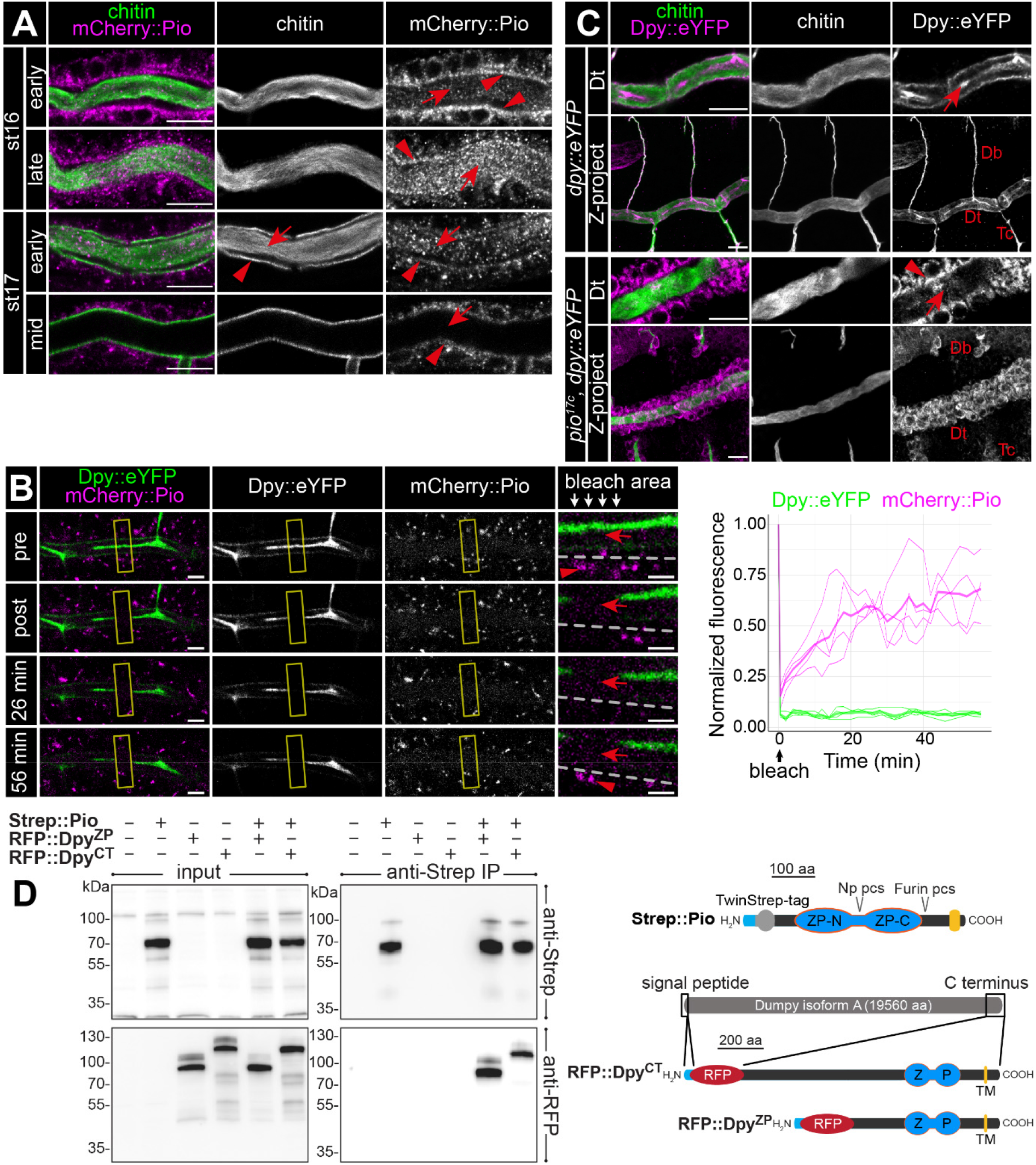
Pio is dynamically localized at the membrane and controls Dumpy secretion. **(A)** Confocal LSM images of dorsal trunks of embryos with endogenous mCherry:: Pio expression stained with anti mCehrry antibody (magenta) and cbp (chitin; green) at indicated embryonic stages. In early stage 16 embryos, mCherry:: Pio enriches apically (arrowheads) and is present in the lumen (arrow). In contrast, at the end of stage 16 mCherry::Pio predominantly localizes within the tracheal lumen (arrows). The luminal mCherry:: Pio staining disappeared during stage 17. **(B)** Confocal images of a representative FRAP experiment (n=4) in a live embryo with endogenous expression of Dpy::eYFP and mCherry::Pio and quantification of normalized fluorescence in the bleached area of n=4 embryos (right) are shown. Yellow frames indicate the bleached area. Close ups (right-most images) show details of the bleached area (below arrows in header). The dashed line indicates apical cell membranes, red arrows indicate luminal mCherry::Pio and red arrowheads indicate intracellular or membrane-associated mCherry::Pio. A representative movie of a FRAP experiment is presented in Movie S2. The mCherry::Pio (magenta) reveals fast recovery of small Pio puncta in the bleached area within the tracheal lumen, while Dpy::eYFP (green) shows no recovery even after 56 min. Scale bars indicate 5 µm in overview panels and 2 µm in bleach close-ups. **(C)** Confocal LSM images of endogenous expression of Dumpy:eYFP stained with anti-GFP antibody. The *wt*-like stage 16 control embryos show extracellular Dumpy:eYFP (magenta) in the aECM at the cell surface and in the luminal cable (arrow) overlapping with cbp (chitin; green). In contrast, in *pio* mutant embryos Dumpy::eYFP did not overlap with chitin (cbp, green), but remained intracellularly (arrowhead). Upper rows focus on the dorsal trunk, lower rows show 3D maximum intensity projections of whole tracheal segments. Note the dorsal branch disruption known from hypomorphic *pio* point mutation allele (Jaźwińska et al., 2003). Single channels are indicated in grey. Db, dorsal branch; Dt, dorsal trunk; Tc, transverse connective. Scale bars indicate 10µm. **(D)** Immunoblotting of co-immunoprecipitation (Co-IP) assay of RFP-tagged Dpy constructs and Strep-tagged Pio expressed in Drosophila S2R+ (Schneider) cells reveals binding of Dpy and Pio. Schemata of expressed proteins used in the assay are shown on the right. Strep::Pio is the full- length Pio protein with a Twin-Strep tag inserted C-terminal to the signal peptide (light blue). RFP::Dpy^ZP^ and RFP::Dpy^CT^ both contain the endogenous Dpy signal peptide (light blue) followed by mCherry (RFP) and different length of the C-terminal region of the Dpy isoform A protein as indicated. Transmembrane (TM) domains (yellow), ZP domains (blue) and Furin and Np protease cleavage sites (pcs) in Pio are indicated. Western blots of input cell lysates (left) and anti-Strep IP elutions (right) stained with anti-Strep (top) and anti-RFP (bottom) antibodies are shown. Both RFP::Dpy proteins are only detectable in IP elutions when they were Co-expressed with Strep::Pio.

### Pio binds Dumpy to organize the luminal ZP protein matrix

Dumpy (Dpy) is a giant (3.2 mDa) and stretchable ZP domain protein. In stage 16 embryos *Dpy::eYFP* (Lye et al., 2014) appears at the tracheal apical cell surface and predominantly within the lumen (Fig. 4C) (Jaźwińska et al., 2003; Dong et al., 2014). In contrast, Dpy::eYFP signal predominantly remained intracellularly in *pio* mutants (Fig. 4C), showing that Dpy secretion depends on Pio. We also performed cell culture experiments to extend our analysis of Dpy secretion. We generated constructs of RFP-tagged Dpy that lacked a portion of EGF and DPY repeats but contained the essential Dpy C-terminal region (ZPD domain, transmembrane domain, cytoplasmic region) (Fig. 4D). Only the co-expression of RFP-tagged Dpy with FLAG-tagged Pio resulted in extracellular RFP::Dpy localization in the S2R+ cells (Fig. S6A). Since extracellular RFP::Dpy is not released from the cells but overlaps with FLAG::Pio at the membrane, it suggests that they co-localize at the cell surface. The S2R+cells expression products of RFP::Dpy constructs were only pulled down together with Strep::Pio in Strep-IP samples (Fig 4D). These data demonstrate that Pio co-localizes and interacts with Dpy. In contrast to Pio, tracheal Dpy::YFP was immobile in our FRAP experiments (Fig. 4B; Video S2), supporting findings that the exchange of Dpy is negligible (Dong et al., 2014). Importantly, the current model suggests that tracheal Dpy-containing matrix stretches very likely by force applied from the cells during tube expansion and thus must be connected to the epithelium (Dong et al., 2014). Our FRAP data suggest that Pio is the dynamic part of the tracheal ZP-matrix, while elastic Dpy modulates mechanical tension within the matrix (Wilkin et al., 2000; Dong et al., 2014).

### Pio release involves the serine protease Notopleural at the apical cell membrane

Furin-like enzymes cleave ZP precursors (Jovine et al., 2005). Pio contains a Furin **p**roteolytic **c**leavage **s**ite (Furin pcs) followed by a C-terminal transmembrane domain. Surprisingly, we detected in stage 16 embryo lysate three mCherry::Pio variants, one correlating with the predicted mass of a full-length protein (115 kDa), a second after furin site cleavage (90kDa), and a third one correlating with the size after proteolysis within the ZP domain (60 kDa) (Fig. 5A). The latter was even the predominant variant in stage 17 (Fig. 5A). In contrast, the Np mutant embryo lysates of stage 16 and 17 contained only faint amounts of the 60 kDa mCherry::Pio and an increased level of the small variant was not apparent in stage 17 embryos (Fig. 5A). These findings support our recent study suggesting that Np cleaves the Pio ZP domain *in vitro* and *in vivo* (Drees et al., 2019). Co-expression of the catalytically inactive Np^S990A^ with mCherry::Pio in *Drosophila* Schneider cells showed as a prominent signal the 90kDa mCherry::Pio variant in the cell lysate (Fig. 5B), and live imaging revealed mCherry::Pio localization at the cell surface (Fig. S6B). This was comparable with control cells that expressed mCherry::Pio alone (Fig. 5B, Fig. S6A). In contrast, cells that co-expressed either the functional Np or its homolog, the human Matriptase, revealed as a predominant signal the 60kDa mCherry::Pio variant in the supernatant fraction (Fig. 5B) and no mCherry::Pio localization at the cell surface (Fig. S6B). These results demonstrate that the enzymatic activity of the serine protease Np is sufficient for ZP domain cleavage, which results in the ectodomain shedding of Pio in Schneider cells.

**Figure 5.**
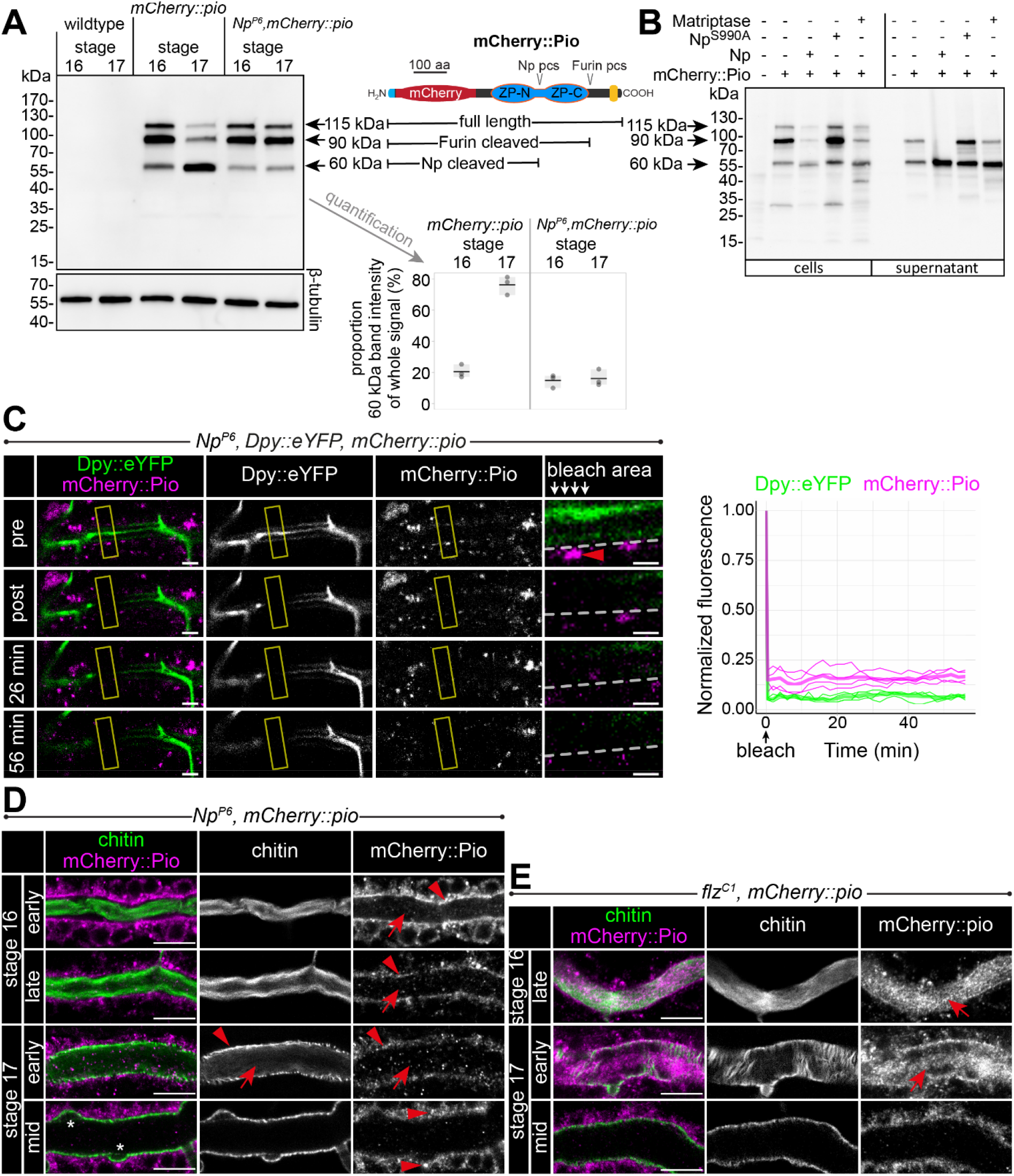
Np-mediated ZP domain shedding controls Pio dynamics at the apical cell membrane. **(A)** Left: Immunoblot of protein lysates from stage 16 and stage 17 embryos stained with anti- mCherry antibody show three specific bands in samples from mCherry::Pio expressing embryos. Middle: Schematic presentation of the mCherry::Pio fusion protein with signal peptide (light blue), mCherry (red), ZP domain (blue) and transmembrane domain (yellow). Furin and Np protease cleavage sites (pcs) and expected molecular weights of resulting fragments are indicated. Right: Proportional intensity of the Np-cleaved 60 kDa mCherry::Pio fragment to the whole signal from all three mCherry::Pio fragments normalized to β-Tubulin intensity. Data from three biological replicates show that proteolytic processing of mCherry::Pio at the Np cleavage site is highly increased in stage 17 embryos compared to stage 16 embryos in *wt* genetic background. This difference is not detectable in samples from *Np* mutant embryos. **(B)** Cleavage assay within the Pio ZP domain is mediated by proteolytic activity of Np and the human Matriptase. Immunoblotting of cell lysates and supernatant precipitates from *Drosophila* S2R+ cells expressing mCherry::Pio alone or together with Np, catalytically inactive Np^S990A^ or human Matriptase with anti-mCherry antibody. Np and human Matriptase cleave mCherry::pio, causing shedding of the mCherry::Pio extracellular domain and a substantial increase of the 60 kDa mCherry::Pio, which correlates in size with cleavage at the ZP domain, in cell culture supernatants. This effect is not observable for catalytically inactive Np^S990A^. **(C)** Confocal images of a representative FRAP experiment in a live *Np* mutant embryo with endogenous expression of Dpy::eYFP and mCherry::Pio and quantification of normalized fluorescence in the bleached area (yellow frames) of n=4 embryos (right) are shown. Close-ups (right-most images) show details of the bleached area (below arrows in header). The dashed line indicates apical cell membranes, arrowheads indicate intracellular or membrane-associated mCherry::Pio. Arrowhead points mCherry::Pio at the apical cell surface in the untreated area. Representative movie of the FRAP experiments is presented in Video S4 (*Np* mutant), compare with Video 2 (*wt*). The fast recovery of small mCherry::Pio puncta in the tracheal lumen is impeded in *Np* mutant embryos (compare with *wt* in Fig. 4D). As in *wt* embryos, Dpy::eYFP (green) shows no recovery even after 56 min. Scale bars indicate 5 µm in overview panels and 2 µm in bleach close-ups. **(D)** Confocal images of tracheal dorsal trunks of *Np* mutant embryos with endogenous expression of mCherry::Pio at indicated developmental stages stained with cbp (chitin; green) and anti- mCherry antibody (magenta). Single channels are indicated in grey. Stage 16 *Np* mutant embryos show intracellular mCherry::Pio at the apical cell surface (arrowhead), which is similar to *wt* control embryos (see Fig. 4A). In contrast to control embryos, the luminal mCherry::Pio (arrow) is strongly reduced in stages 16 and 17 *Np* mutant embryos, while the non-luminal mCherry::Pio accumulates in stage 17. The luminal chitin cable is degraded normally in *Np* mutant embryos but does not condense (Drees et al., 2019) during early stage 17 and instead, remains attached to the tracheal cell surface and fills the whole lumen during degradation (compare with *wt* in Fig 4A and Fig 6A). The asterisks mark bulges in *Np* mutant tubes. Note the two layers of chitin visible at the membrane bulges and the adjacent aECM indicating disintegration of the tracheal chitinous aECM (see also Fig. 6). Scale bars indicate 10 µm. **(E)** The tracheal trypsin-like S1A Serine transmembrane protease Filzig (Flz) shows high sequence homology to Np (Drees et al., 2019) and acts in processing the lumen matrix (Rosa et al., 2018). Confocal images of dorsal trunks of *flz* mutant embryos with endogenous expression of mCherry::Pio at indicated developmental stages stained with cbp (chitin; green) and anti mCherry antibody (magenta). Single channels are indicated in grey. The *flz* mutant embryos revealed normal Pio expression, luminal shedding and clearance from airways (compare with *wt* in Fig 4A). In contrast to *Np* mutant embryos, the luminal aECM cable condensed during early stage 17 luminal clearance (arrows) as in *wt* embryos (compare with *Np* mutant in D and *wt* in Fig 4A and Fig 6A). Scale bars indicate 10 µm.

Next, we investigated the consequences of NP-mediated ZP cleavage. FRAP experiments showed only minor intracellular recovery of mCherry::Pio in *Np* null mutant embryos (Video S4). In contrast to the control, extracellular mCherry::Pio recovery within the tube lumen failed 56 min after bleaching in *Np* mutant embryos (Fig. 5C, Video S3). The luminal mCherry::Pio immobility was comparable to the immobile Dpy::eYFP fraction in control and *Np* mutant embryos, suggesting that Pio is hampered to diffuse in *Np* mutant embryos. Second, mCherry::Pio was not released into the tracheal lumen in stainings of late-stage 16 *Np* mutant embryos (Fig. 5D, Video S4). Third, the tracheal trypsin-like S1A Serine transmembrane protease Filzig had no influence on the mCherry::Pio pattern (Fig. 5E, Fig. S7). The *flz* mutant (Fig. S7E) and *Np* mutants st17 embryos showed no Dpy clearance from the tracheal lumen (Drees et al., 2019). However, the *flz* mutation did not affect the release of Pio at the apical cell surface. Further, the luminal aECM cable condensed during early stage 17 in *flz* mutant embryos as observed in wt (compare Fig. 4A with Fig. 5D,E), indicating that these processes require Np. In summary, our findings prove the inhibition of luminal Pio release in *Np* mutant embryos. Furthermore, the Np-mediated proteolysis of the Pio ZP domain is specific and plays a central role in Pio dynamics at and near the apical cell membrane during tube expansion.

To further analyze Np function, we used Uif to examine the tracheal apical cell membrane structure. This revealed unusual bulge-like membrane deformations in late stage 16 and 17 Np mutant embryos, while control embryos did not show such bulges (Fig. 6A). Confocal time-lapse series revealed that bulge-like deformations emerged during early stage 16 embryos and grew in size (Fig. 6B). Furthermore, the increasing size of the membrane bulges led to the detachment of α- tubulin::GFP marked cells from the aECM at the cell surface (Fig. 6B). Such a detachment between cells and adjacent aECM was not detected in *wt* control embryos (Fig. 1B). However, confocal images of Uif and chitin show residual chitin at the Uif marked membrane and chitin at the detached aECM (Fig. 6A). These results indicate the tearing of the tracheal aECM at the apical cell surface.

**Figure 6:**
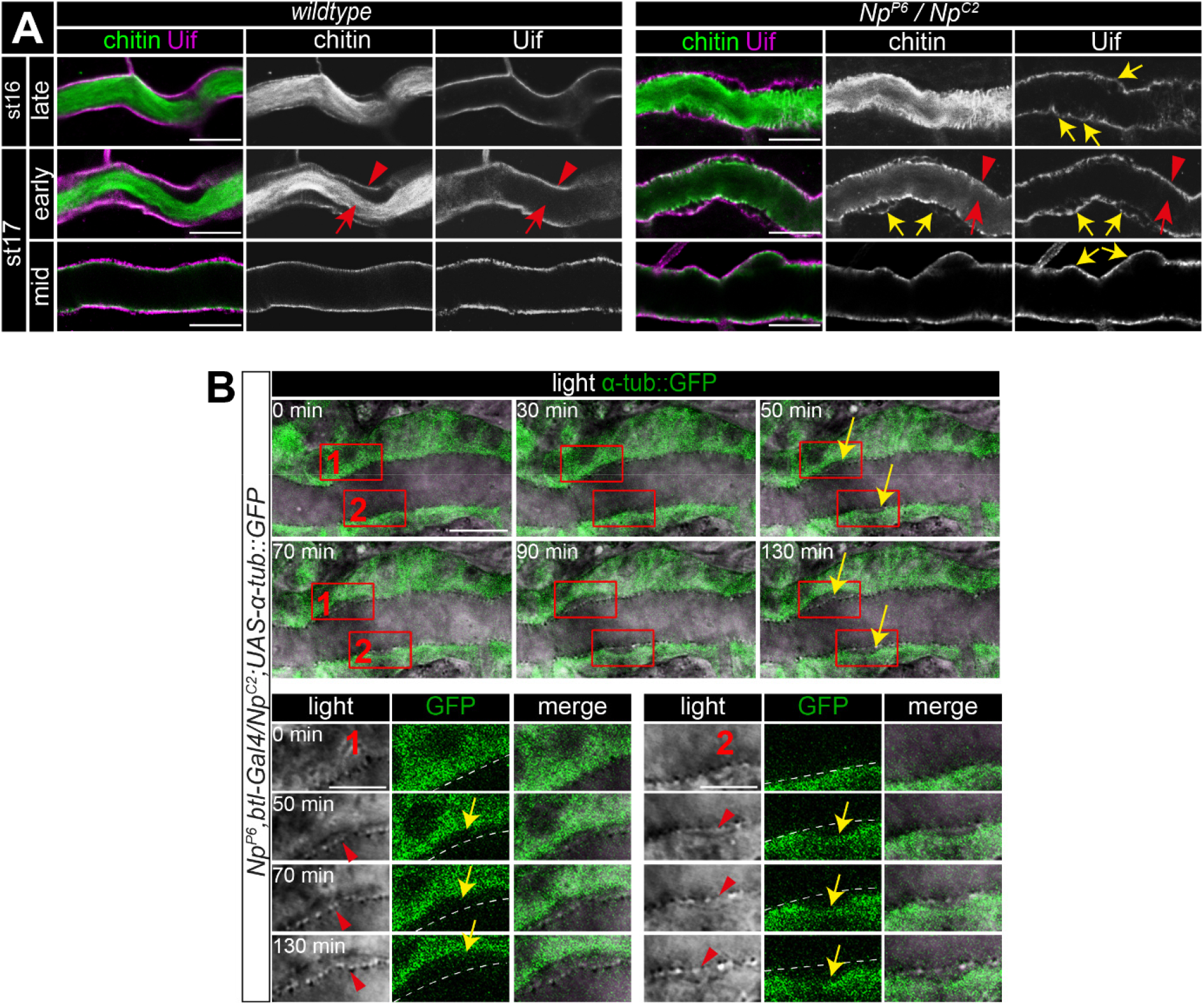
Np supports structural cell membrane integrity. **(A, B)** Bulge-like tracheal apical membrane deformations appeared in *Np* mutant embryos as stable structures that grow during late stage 16. Confocal images of dorsal trunks of *wt* embryos and *Np^P6^ / Np^C2^* embryos stained with cbp (chitin; green) and anti-Uif antibody (magenta) as a marker for apical tracheal membranes (A). The *in vivo* time laps series of 130 min show bulges arising at the dorsal trunk cell membranes of Np mutant embryos (B). Tracheal cells express Tubulin::GFP in transheterozygous *Np^P6^;Np^C2^* mutant embryos. Frames 1 and 2 are shown as close-ups (below) of forming bulges. The cell membrane of GFP expressing cells and parts of the misorganized tracheal cuticle aECM (dashed line) separate during the time-lapse. Yellow arrows point to membrane deformations; red arrowheads point to tracheal cuticle at the apical cell surface and red arrows to the luminal aECM cable. Note that chitin is detectable in two layers at the sites of bulges, while Uif is detectable only at the bulges, indicating a disintegration of the tracheal chitinous aECM (A). Scale bars are 10 µm (overviews) and 3.5 µm (details). Note that bulges grew in size as time progressed (up to 130 min).

Finally, our time laps series and the analysis of stage 17 embryos proved that the bulge-like cell membrane structures remained in *Np* mutant embryos. Moreover, these cell membrane bulges destabilize the normal epithelial barrier. Upon tracheal expression of myr-RFP in Np mutants, we detected accumulation of RFP signal first at the membrane bulge and subsequently in the tube lumen (Fig. S8). The *Np* and *pio* mutant embryos show apicel membrane deformations (compare Figs. 1A and 6A). Np is located at the membrane (Drees et al., 2019), where it controls Pio release into the tracheal lumen. This suggests that membrane deformations in *Np* mutant embryos are caused by defective function of the Pio-mediated ZP matrix due to a lack of Pio shedding.

Any imbalance between membrane and matrix during tube expansion causes tube deformations. Chitin staining was reduced and revealed sinusoidal over-elongated tubes in stage 16 *pio* mutant embryos (Fig. 7A,B), proving that *pio* function prevents tracheal tubes from over-elongation. In *Np* mutant stage 16 embryos, Pio was not shedded into the lumen but remained at the cell, while Dumpy showed normal and immobile localization within the tube lumen (Fig. 5C,D; Video S3,S4). This indicates that the Pio-mediated ZP matrix can still restrict tube expansion of the membrane in Np mutants, independently from membrane deformations. The *Np* mutant stage 16 embryos showed normal tube sizes (Fig. 7A,B). Further, *Np,pio* double mutants did not exacerbate the *pio* mutant tube length defects suggesting that both act in the same genetic pathway (Fig. 7A,B). Our data assumes that Np overexpression may enhance Pio shedding affecting the Pio-mediated ZP matrix function. Consistently tracheal Np overexpression led to tube overexpansion in stage 16 embryos resembling the *pio* mutant phenotype (Fig. 7A,B). Thus, Np-mediated Pio shedding controls Pio function.

**Figure 7:**
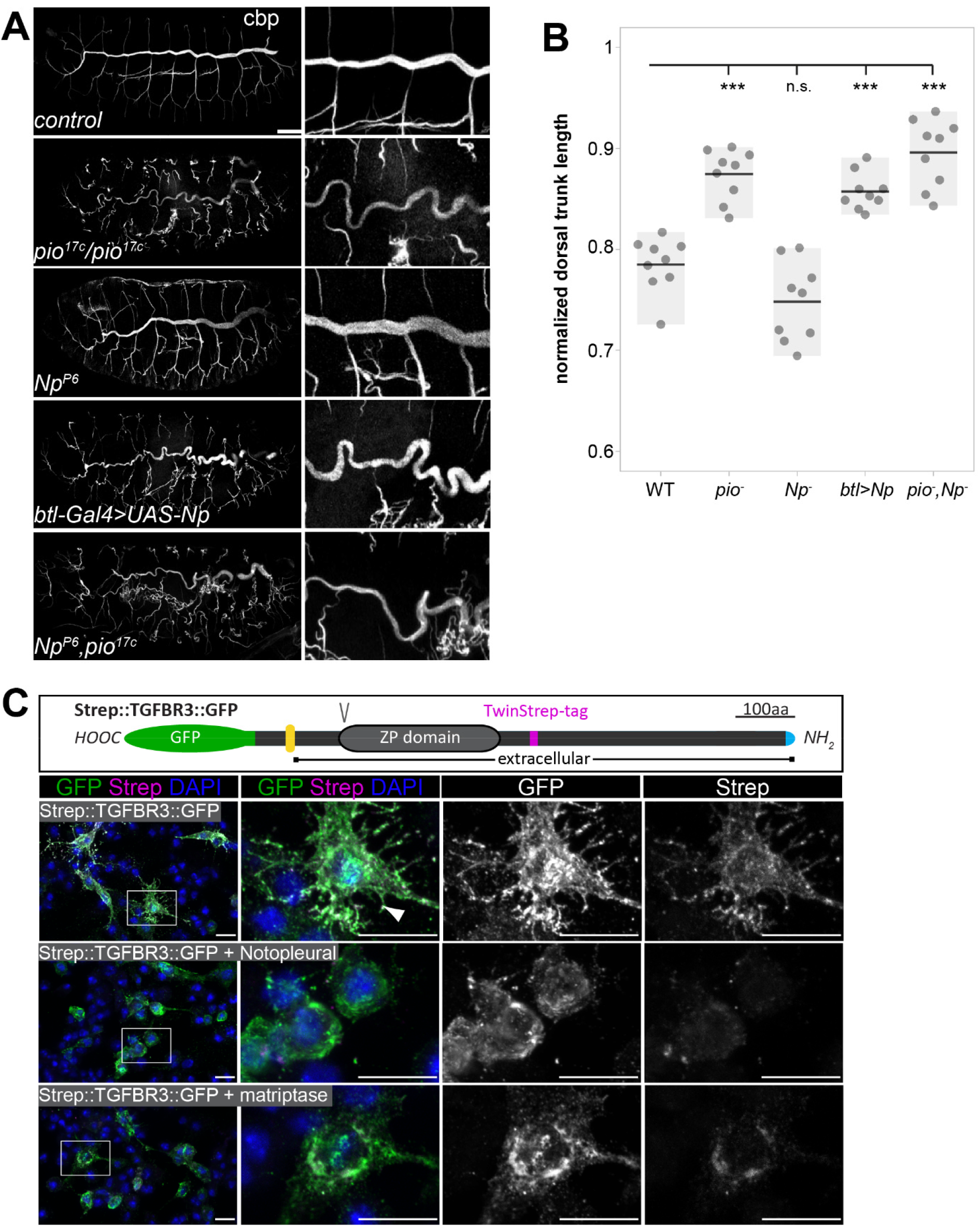
Pio and Np control tube size and their regulatory mechanisms of ZP domain shedding is conserved. **(A)** Confocal Z-stack projections of whole mount stage 16 embryos stained with cbp (chitin) focusing on the tracheal system and close ups at the right. In contrast to straight branches of control embryos, *pio^17C^* null mutant embryos revealed curly elongated tubes indicating excess tube expansion. Note dorsal branch disruption known from a hypomorphic *pio* point mutation (Jaźwińska et al., 2003). *Np* mutant embryos show straight *wt*-like tubes. Embryos that overexpress *Np* in the tracheal system (*btl-Gal4>UAS-Np*) show curly elongated tubes and dorsal branch disruption, phenocopying the *pio* mutant phenotype. *Np;pio* double mutant embryos do not exacerbate *pio* mutant tube size defects and show a similar phenotype as *pio* mutant embryos respectively. **(B)** Quantification of normalized dorsal trunk length from 9 stage 16 embryos of each genotype. Anterior-posterior dorsal trunk length was divided by the anterior-posterior length of the embryo. Normalized dorsal trunk length in *pio* mutant embryos, *btl-Gal4>UAS-Np* embryos and *Np*,*pio* double mutant embryos is significantly increased when compared with *wt* (p=0.00013, p=0.00007, p=0.00019). Notably, the Np mutant dorsal trunk is relatively straight, while control embryos show slightly convoluted tubes. Also statistical analysis reveals the tendency of slightly shortened dorsal trunk length in Np mutant. Individual points represent the respective embryos. **(C)** Human TGFβ type III receptor (TGFβRIII) is a widely expressed ZP-domain containing protein. Human TGFβRIII with a cytoplasmic GFP tag and an extracellular Strep tag was expressed in *Drosophila* S2R+ cells either alone or together with *Drosophila* Np or human Matriptase. A schema of the tagged TGFβRIII is shown (top). The ZP and transmembrane (yellow) domains, the N- terminal Strep tag (magenta), C-terminal GFP (green) Furin protein cleavage site (V) and the signal peptide (blue) are indicated. Images display maximum intensity projections of confocal Z- stacks. Shown are S2R+ cells that expressed the Strep::TGFβRIII::GFP construct alone or together with Np or human Matriptase stained with DAPI (blue) anti GFP (green) and anti-Strep (magenta) antibodies. Single channel panels are indicated in grey. Control cells contain co- localizing GFP and Strep signals. The co-expression of Np or Matriptase reveals strong GFP but faint Strep signals due to extracellular cleavage and shedding of the TGFβRIII ectodomains. Framed boxes in overview images display details in panels on the right side. Scale bars indicate 10 µm.

Since our findings show that Np controls tracheal Pio function by ZP domain cleavage, we addressed whether this is also a putative mechanism of human ZP domain proteins required in the lung. Idiopathic pulmonary fibrosis (IPF) is associated with a progressive loss of lung function and is characterized by fibroblast accumulation and relentless deposition of extracellular matrix (ECM) (King et al., 2011; Loomis-King et al., 2013). Upregulation of matriptase activity in human IPF cells and the corresponding experimental mouse model promotes pulmonary fibrogenesis (Bardou et al., 2016). Furthermore, the lungs of IPF patients and mouse model have increased Transforming growth factor-β receptor density on their surface (Naik et al., 2012). The type III Transforming growth factor-β receptor (TβRIII) acts as a signaling modifier and co-receptor of TGF-β and contains a ZP domain (López-Casillas et al., 1991; Moustakas et al., 1993). TβRIII- ectodomain shedding in lung cancer cell models induces epithelial-to-mesenchymal transition and promotes growth of tumors (Huang et al., 2019). For an *in vitro* cleavage assay, we co-expressed GFP- and Strep-tagged combined variants of TβRIII together with either human Matriptase, or Np in *Drosophila* cells. The C-terminal GFP-tag detects the intracellular part of TβRIII. The N-terminal Strep-tag follows the extracellular ZP domain and recognizes the ectodomain. When cells expressed TGFβRIII without Matriptase or Np they revealed strong colocalization signals of GFP and Strep. Co-expression of either human Matriptase or Np caused a substantial reduction of the extracellular Strep signal while the intracellular GFP signal remained (Fig. 7C). This shows that matriptases catalyze the extracellular proteolysis and thereby the extracellular localization of the human TβRIII ZP domain protein. This indicates a new and unexpected conserved mechanism with the capability to control TβRIII function in the lung and other epithelia.

## Discussion

Tracheal tube lumen expansion requires mechanical stress regulation at apical cell membranes and attached aECM. This involves the proteolytic processing of proteins that set local membrane-matrix linkages. Thus, the membrane microenvironment exhibits critical roles in regulating tube and network functionality.

ZP domain proteins organize protective aECM in kidney, tectorial inner ear, and zona pellucida (Jovine et al., 2005; Litscher and Wassarman, 2020), as well as in *Drosophila* epidermis, tendon cells and appendages (Bökel et al., 2005; Plaza et al., 2010; Ray et al., 2015). *Drosophila* ZP domain proteins link the aECM to actin and polarity complexes in epithelial cells (Fernandes et al., 2010). Dumpy establishes force-resistant filaments for anchoring tendon cells to the pupal cuticle (Chu and Hayashi, 2021).

We identify that ZP protein-mediated microenvironmental changes increase the flexibility of membrane-matrix association, resulting from the activity of ZP domain proteins (Figs. 1,6). Shear stress stimulates the activity of membrane-anchored proteases (Kang et al., 2015) and potentially also Np since we did not observe the misdistribution of the tracheal cytoskeleton when blisters arise. The dynamic membrane-matrix association control is based on our findings that loss of Np prevents Pio ectodomain shedding at the apical cell membrane resulting in immobile localization of Pio at the membrane and Dpy localization within the matrix (Fig. 5). Direct interaction and overlapping subcellular localization at the cell surface showed that both proteins form a ZP matrix that potentially attaches membrane and ECM (Figs. 2,4). Deregulation of Pio shedding blocks ZP matrix rearrangement and release of membrane-matrix linkages under tube expansion and subsequent shear stress. This destabilizes the microenvironment of membranes, causing blister formation at the membrane due to ongoing membrane expansion (Fig 6). Additionally, Pio could be part of a force-sensing signal transduction system destabilizing the membrane and matrix. Our observation that the membrane deformations are maintained in *Np* mutant embryos supports our postulated Np function to redistribute and deregulate membrane-matrix associations in stage 16 embryos when tracheal tube length expands. In contrast, Np overexpression potentially uncouples the Pio-Dpy ZP matrix membrane linkages resulting very likely in unbalanced forces causing sinusoidal tubes (Fig 7).

The membrane defects observed in both Pio and Np mutants indicate errors in the coupling of the membrane matrix due to the involvement of Pio (Figs. 1,6). In *pio* mutants, gaps appear between the deformed membrane and the apical matrix (Fig. 1B-D). These changes in apical cell membrane shape are consistent with increased cell and tube elongation in *pio* mutant embryos because the matrix is uncoupled from the membrane in such mutants (see model Fig. S9). In contrast to *pio* mutants, the large membrane bulges in *Np* mutants affect the membrane and the apical matrix (Fig. 6). Since apical Pio is not cleaved in *Np* mutants (Fig. 5D), the matrix is not uncoupled from the membrane as in *pio* mutant embryos but is likely more intensely coupled, which leads to tearing of the matrix axially along the membrane bulges (Fig. 6), when the tube expands in length (Fig. S9). If apical Pio detachment reduces coupling between the matrix and apical membrane, then it is likely that *Np* mutant embryos may exhibit a reduced tube length phenotype. Indeed, in Np mutant embryos, tracheal dorsal trunk length is slightly reduced compared to *wt* embryos (Fig. 7B), suggesting that Pio shedding plays a critical role in controlling tracheal tube lumen length. We assume that Np overexpression increases Pio shedding, resulting in a *pio* loss-of-function phenotype. Thus, the tube length overelongation upon Np overexpression indicates that non-shedded apical Pio is required for tube length control.

Is Pio ectodomain shedding in response to tension? We did not measure tension directly. However, the developmental profile of mechanical tension during tracheal tube length elongation in stage 16 embryos (Dong et al., 2014) is consistent with the profile of Pio shedding. Apical Pio is predominantly cleaved by Np during stage 16 when tube length expands (Fig. 4A). In contrast, Pio shedding decreases sharply at early stage 17 when tube elongation is completed. Our model, therefore, predicts that loss of Pio or increased Pio secretion at stage 16 may reduce the coupling of the membrane matrix so that increased tracheal tube elongation is maintained until the end of stage 16, which is found in *pio* mutants and upon Np overexpression (Fig. 7A,B). Additional unknown mechanisms, such as distinct membrane connections during development and emerging connections to the developing cuticle, may also influence tension at the apical membrane during tube length control.

Our studies on human matriptase provide evidence for the mechanistic conservation of ZP-domain protein as a substrate for ectodomain shedding (Fig. 7). Matriptase degrades receptors and ECM in pulmonary fibrinogenesis in squamous cell carcinoma (Bardou et al., 2016; Martin and List, 2019). TβRIII is a membrane-bound proteoglycan that generates a soluble form upon shedding (López-Casillas et al., 1991), a potent neutralizing agent of TGF-β. Expression of the soluble TβRIII inhibits tumor growth due to the inhibition of angiogenesis (Bandyopadhyay et al., 2002). Idiopathic pulmonary fibrosis (IPF) is associated with a progressive loss of lung function due to fibroblast accumulation and relentless ECM deposition (King et al., 2011; Loomis-King et al., 2013). Upregulation of matriptase activity and increased TGF-β receptor density affect human and mouse model IPF cells on pulmonary fibrogenesis (Naik et al., 2012; Bardou et al., 2016). Interestingly, human Matriptase induces the release of proinflammatory cytokines in endothelial cells, which contribute to atherosclerosis and probably also to abdominal aortic aneurysms (Seitz et al., 2007). Membrane bulges arising in our *Drosophila* model during tracheal tube elongation upon Np loss of function showed analogy to the appearance of artery aneurysms. Bulges with varying phenotypic expression in different organs can lead to aortic rupture due to fragile artery walls or degeneration of layers in responses to stimuli, such as shear stresses (Kubo et al., 2015). Indeed, aneurysms development is forced by alterations in the ECM (Yoon et al., 1999) and are characterized by extensive ECM fragmentation caused by shedding of membrane-bound proteins (Yoon et al., 1999; Antalis et al., 2016; Quintana and Taylor, 2019; Zhong and Khalil, 2019).

We identified a dynamic control of matrix proteolysis, very likely enabling fast and site-specific uncoupling of membrane-matrix linkages when tubes expand. Such a scenario has not yet been studied in angiogenesis. It may represent a new starting point for genetic studies to decipher the putative roles of ZP domain proteins and matriptase in clinically relevant lung syndromes, including the formation of aneurysms and IPF caused by membrane deformation and defects in size determination of airways and vessels.

## Materials and Methods

### Fly husbandry, gas-filling, and statistics

For collection of *D. melanogaster* embryos and larvae, flies of the desired genotype took place at 25 °C for collection of embryos for RNAi-mediated knockdown. All used fly strains are listed in the Materials and Methods section in the supplement. In all other cases, egg-laying took place at 22 °C. The apple-juice agar plates were exchanged with fresh plates at respective points of time to obtain embryos or larvae at certain developmental stages. The mutant alleles were kept with “green” balancers to recognize mutant embryos.

We used w^1118^ as control (referred to as wild-type, *wt*). Rescue experiments: *w*; pio17C, btl-Gal4 /Cyo, dfd-eYFP* were mated with *w*; pio5M /Cyo, dfd-eYFP ; UAS-Pio* to receive the rescue in the progenoty *w*; pio17C, btl-Gal4 / pio5M ; UAS-pio / +.* For gas filling assay, we transferred stage 17 embryos and freshly hatched larvae onto agar plates and studied those by bright field microscopy. Significance was tested using t-tests in Excel 2019; asterisks indicate p-values (* p<0.05, ** p<0.01, ***p<0.001); error bars indicate the standard deviation.

### Embryo dechorionation, fixation and immunostainings

In general, we analyzed for control minimum n>20 embryos, and for *pio* or other mutants n>10 embryos. Embryos were washed from the apple-juice agar plates into close-meshed nets, incubated for 3min in a bleach solution (2,5% Sodium hypochlorite) for dechorionation.

For subsequent fixation, embryos were incubated at 250 rpm for 20 min in 1 ml 10 % (v/v) Formaldehyde solution (50 mM EGTA, pH7,0), 2 ml Hepes solution and 6 ml heptane. The fixative was removed and 8 ml methanol added and incubated at 500 rpm for 3 min to detach the vitelline membrane. Finally, embryos were washed with methanol and stored at -20 °C.

All used antibodies are listed in the Materials and Methods section in the supplement. For antibody staining, embryos were 5min washed three times with BBT followed by blocking for 30 min. Subsequently, pre-absorbed primary antibodies diluted in blocking solution were applied to the embryos and incubated over night at 4 °C. Primary antibody was incubated overnight washed off six times with BBT. After blocking for 30 min pre-absorbed secondary antibodies diluted in blocking solution were added to the embryos and incubated for 2 hrs. If required, Alexa488-conjugated Chitin-binding-probe (cbp) was added to the secondary antibody dilutions at a 1:200 dilution for staining of chitin. Finally, embryos were washed six times with PBT for 5 minutes and mounted either in Prolong mounting medium (Thermo Fisher Scientific) or in phenol free Kaiser’s glycerol- gelantine (Carl Roth).

### Light microscopy and image acquisition

For bright field and dark field light microscopy we used an Axiophot microscope (Zeiss; A-PLAN 10x/0.25) and images were acquired with an AxioCam (HRc / mRc) and the Zen acquisition software (Zeiss). For handling confocal analysis, we used ZEN software (2.3, SP1 black) and Zeiss LSM780-Airysacn (Zeiss, Jena) microscopes. Overview imaging were taken with 25x/0.8 LD-PLAN Neofluar and magnifications with Plan-Apochromat 40x/1.4 Oil DIC M27 and with 63x/1.3 PLAN Neofluar M27 objectives and Zeiss oil/water/glycerol medium. For imaging standard ZEN confocal microscopy (Pinhole Airy1) settings and image processing (maximum intensity projection and orthogonal section) and orthogonal projections were used. The “express” deconvolution of SVI Huygens pro was used with standard settings. Images were transferred to ZEN for further analyses. For Airyscan acquisition, standard ZEN settings and optimal Z-stack distances were used. For 3D visualization, deconvolved confocal and Airyscan Z-stacks were processed in Imaris (version 9.7.2) to convert voxel-based data into surface objects. We cropped Images with Adobe CS6 Photoshop and designed figures with CS6 Illustrator.

### CRISPR/Cas9-mediated mutagenesis and genome editing

Using CRISPR/Cas9 mediated mutagenesis, we generated three independent *pio* lack of function alleles [*pio^17C^*, *pio^5M,^* and *pio^11C^*] and one *flz* lack of function allele (*flz^C1^*) that carry frameshift mutations in the ORF. The *pio* mutations led to truncated Pio proteins containing only short N-terminal stretches but lacking all critical for Pio function. These mutations in *pio^17C^*, *pio^5M^* and *pio^11C^* alleles and *flz^C1^* allele caused embryonic lethality. The *in vivo* analysis of airways in *pio^17C^*, *pio^5M,^* and *pio^11C^* homozygous and *pio^17C^*/*pio^5M^* transheterozygous mutant stage 17 embryos revealed lack of tracheal air-filling. The sequences of sgRNAs are listed in the Materials and Methods section in the supplement.

The CRISPR Optimal Target Finder tool was used to identify specific sgRNA targets in the *D. melanogaster* genome. DNA oligos corresponding to the chosen sgRNA target sequences were annealed and ligated into the *pBFv-U6.2* vector via BbsI restriction sites (Kondo and Ueda, 2013). For simultaneously targeting two sgRNA recognition sites, one sgRNA target encoding annealed DNA oligo was ligated into the *pBFv-U6.2* vector and the other one into the *pBFv-U6.2B* vector. Subsequently, the *U6.2* promotor and sgRNA target sequence of the *pBFv-U6.2* vector were transferred to the sgRNA target sequence containing *pBFv-U6.2B* vector via EcoRI/NotI endonuclease restriction sites. Vectors for sgRNA expression were injected into *nos-Cas9-3A* embryos by BestGene. Adult flies that developed from the injected embryos were then crossed with balancer chromosome strains. Single flies of the resulting F1 generation were again crossed with balancer chromosome strains and the resulting F2 stock was analyzed regarding lethality of the putatively mutated chromosome. The regions of the genomic sgRNA target sites from the obtained lethal stocks were amplified by PCR followed by purification of the amplicon and sequencing.

The CRISPR/Cas9 system was used for targeted insertion of transgenes by homology-directed repair. Sequences of homology arms (HAs) were amplified by PCR from genomic DNA of *nos- Cas9-3A* flies. HAs were then purified and cloned into the *pHD-ScarlessDsRed* donor vector for editing of *pio*. A mCherry encoding sequence was inserted into the *pio* ORF at the 3’ end of the 5’ HA in the *pHD-ScarlessDsRed* donor vector. The donor vector together with the vector for sgRNA expression, were injected into *nos-Cas9-3A* embryos by BestGene. Adult flies that developed from the injected embryos were crossed with *w\** flies and resulting F1 flies were selected for presence of *3xP3-DsRed* marker. Stocks were then established by crosses with balancer chromosome strains. Correct insertion of the transgenes was verified by amplification of the targeted genomic regions by PCR and subsequent purification of the amplicons and sequencing. The *3xP3-DsRed* marker gene that was inserted in the *pio* locus was removed by crosses with the *tub-PBac* strain and subsequent selection of F2 flies that lacked the *tub-PBac* balancer chromosome and the *3xP3-DsRed* marker gene.

### Cell culture-based experiments

*D. melanogaster* S2R+ cells (DGRC) and Kc167 cells (DGRC) were kept in flasks containing Schneider’s *Drosophila* medium (Thermo Fisher Scientific) supplemented with 1 % Penicillin/Streptomycin (Thermo Fisher Scientific) and 10 % FBS (Sigma-Aldrich) at 25 °C. Handling of cells was performed in the sterile environment of a clean bench.

Confluent cells were detached, diluted 1:6, and transferred either to 10 cm diameter petri dishes, or to 6-well plates (Greiner Bio One), or to glass bottom micro-well dishes (MatTek; 6 ml per dish), approximately 24 hours before transfection. The UAS/Gal4 system was used for protein over- expression in cultured cells. Cells in 6-well plates and glass bottom micro-well dishes were transfected with 500 ng *actin5C-Gal4* vector and 500 ng *pUAST*-responder vector (1 µg total DNA), while cells in 10 cm diameter petri dishes were transfected with twice the amount of each vector (2 µg total DNA). The Effectene transfection reagent (Qiagen) was used for cell transfection according to suppliers’ guidelines. For transfections of cells in 6-well plates and glass bottom micro-well dishes, vector DNA was mixed with 190 µl Effectene EC buffer (Qiagen) followed by adding 8 µl Effectene Enhancer (Qiagen), vortexing and incubation for 5 min. Subsequently, 20 µl Effectene were added and the mix was vortexed for 10 sec followed by incubation for 15 min. The transfection mix was then carefully trickled onto the cells with a pipette. Incubation steps were performed at room temperature. For transfections of cells in 10 cm diameter petri dishes with 2 µg DNA, volumes of the transfection mix components were doubled.

Cells in glass bottom micro-well dishes were incubated at 25 °C for approximately 48 hours after transfection followed by imaging with a confocal LSM.

For immunostainings cells were incubated at 25 °C for 48 hours after transfection and then washed with PBS twice and fixed for 15min by adding formaldehyde solution. The cells were then washed twice with PBS and incubated in PBTX for 150 sec for permeabilization. Subsequently, cells were washed three times with PBS followed 30min blocking. Primary antibodies diluted in blocking solution were added to the cells and incubated for 2 hours. Then cells were washed three times with PBS followed by incubation with secondary antibodies diluted in blocking solution for 1 hour. Finally, the cells were washed twice with PBS followed by mounting in Prolong mounting medium with DAPI (Thermo Fisher Scientific). Images were acquired with a confocal LSM. All incubation and washing steps were performed at room temperature.

For pulldown assays Strep-tagged Pio was co-expressed in S2R+cells together with either of two independent RFP tagged Dpy fragments for pull-down assays. One RFP tagged Dpy contains the Dpy C-terminal region, and the second exclusively the Dpy-ZP domain. Expression products of both RFP::Dpy constructs were only pulled down together with Strep::Pio in the Strep-IP samples. For Western detection see below. All used oligos to generate constructs are listed in the Materials and Methods section in the supplement.

### Protein purification and Western blotting

We dechorionated embryo collection, squashed them with a needle and pulsed with ultrasound for 30 sec, added 25ml 4xSDS sample buffer, heated for 9 min at 96°C, centrifuged at 11000rpm for 20 min. Lysate was stored in a fresh cup at -80/-20°C. Schneider’s cells were manually detached, centrifuged at 900rpm, PBS was replaced by 35µl 4x SDS sample buffer and heated 9min at 96°C and stored at -80°C. For supernatant samples, we centrifuged at 15000rpm at 4°C for 20min, washed with acetone and resuspended in 100µl 1x SDS sample buffer, heating it at 96°C for 9min. We used 4-20% gradient Mini-Protean TGX Precast Protein Gels (BioRad) together with PageRuler prestained protein ladder (Thermo Scientific), MiniProtean chamber (BioRad), PowerPac Basic power supply (BioRad) for 40 min at 170 V. The gels were then equilibrated in transfer buffer and packed into a western blotting sandwich. Sandwich Blotting was performed with a PVDF transfer membrane with 0,2 µm pore size (Thermo Fisher Scientific) for embryos lysates or in other cases, Amersham Hybond-ECL membrane with 0,2 µm pore size (GE Healthcare). Protein transfer to the membrane was performed on ice in a MiniProtean chamber (BioRad) that was filled with transfer buffer at 300 mA for 90 min.

### Fluorescence recovery after photobleaching (FRAP)

The stage 16 embryos were dechorionated, transferred and fixed on the cover slip of small petri dishes (ibidi, Germany; https://ibidi.com/dishes/185-glass-bottom-dish-35-mm.html) and mounted in PBS. We used LSM780 confocal for FRAP experiments. To define the Region of interest (ROI) we used the Zeiss Zen software. The bleaching was performed with 405nm full laser power (50mW) at the ROI for 20 seconds. A Z-stack covering the whole depth of the tracheal tubes in the ROI were taken at each imaging step. The confocal images were taken every 2 min until 60 min after bleaching. To correct for movements of the embryos, images that are presented in figures and supplemental movies were manually overlayed to center them in the same focal plane and to correct for movements in the x and y axis. Fluorescence intensity in the bleached ROIs was measured after correction for embryonic movements using Fiji.

### Electron Microscopy

Stage 17 *Drosophila* embryos were dechorionated, transferred to a 150 μm specimen planchette (Engineering Office M. Wohlwend GmbH), and frozen with a Leica EM HBM 100 high-pressure freezer (Leica Microsystems). Vitrified samples were embedded with an Automatic Freeze Substitution Unit (EM AFS; Leica Microsystems) at −90 °C in a solution containing anhydrous acetone, 0.1% tannic acid, and 0.5% glutaraldehyde for 72 h and in anhydrous acetone, 2% OsO4, and 0.5% glutaraldehyde for additional 8 h. Samples were then incubated at −20 °C for 18 h followed by warm-up to 4 °C. After subsequent washing with anhydrous acetone, embedding in Agar 100 (Epon 812 equivalent) was performed at room temperature. Counterstaining of ultrathin sections was done with 1% uranylacetate in methanol.

Alternatively, the dechorionated embryos were fixed by immersion using 2 % glutaraldehyde in 0.1 M cacodylate buffer at pH 7.4 overnight at 4°C. The cuticle of the embryos was opened by a cut to allow penetration of fixative. Postfixation was performed using 1% osmium tetroxide. After pre-embedding staining with 1% uranyl acetate, tissue samples were dehydrated and embedded in Agar 100. Ultrathin sections were evaluated using a Talos L120C transmission electron microscope (Thermo Fisher Scientific, MA, USA)

## Acknowledgments

We are very grateful to Markus Affolter, Bernard Moussian and Stefan Luschnig for sharing generously antibodies. We thank Christian Wolf for critical reading and helpful discussion. The FlyBase, NCBI and SMART internet tools were always important sources. Stocks obtained from the Bloomington Drosophila Stock Center (NIH P40OD018537), the Vienna Drosophila Resource Center (VDRC) and from Drosophila Genomics Resource Center (NIH grant 2P40OD010949) were used in this study. M.B. expresses his gratitude to Johannes Kacza for his support with Imaris. We appreciate the University Leipzig BioImaging Core Facility (BCF equipment INST 268/230-1; INST 268/293-1; SFB-TR67; EFRE 100192650, 100195814, 100144684) for assistance.

## Author contributions

L.D & M.B designed and conducted the experiments; D.R. performed ultrastructure experiments and analysis; All authors discussed all experiments and results; M.B. wrote and revised the manuscript; All authors discussed and reviewed the manuscript.

