## Supplemental data for "The proteolysis of ZP proteins is essential to control cell membrane structure and integrity of developing tubes"

#### Supplemental Information

##### Video S1 and Video S4

Confocal time-lapse movie of endogenously expressed Dpy::eYFP (magenta) and mCherry::Pio (green) in *wt* control (Video S1) and *Np* mutant embryo (Video S4). The video S1 shows tracheal Pio and Dumpy localization in the dorsal trunk from stage 16 until mid-stage 17. The mCherry::Pio dynamic trafficking reveals ongoing secretion, apical localization, and subsequent shedding into the tubular lumen. Dpy::eYFP shows static luminal localization and undergoes degradation during stage 17. In *Np* mutant embryo, mCherry::Pio shows intracellular and apical localization but no luminal shedding. The Dpy::eYFP shows static luminal localization but is not degraded during tracheal maturation. Images were obtained every 10 minutes.

##### Video S2 and Video S3

Representative confocal time-lapse movie of a FRAP experiment in a *wt* control embryo (Video S2) with endogenous expression of Dpy::eYFP (green) and mCherry::Pio (magenta). The bleached area is indicated by a white box. The first frame is before bleaching and subsequent images were obtained every two minutes as indicated in the middle panel. mCherry::Pio shows fast recovery in the tube lumen already after two minutes and steady recovery of intracellular foci during the 56 min time lapse. Luminal Dpy::eYFP shows no recovery in the bleached area. In *Np* mutant embryo (Video S3), mCherry::Pio foci are visible intracellularly as in *wt* (Movie S2), but almost no mCherry::Pio is detectable in the tracheal lumen. After bleaching, mCherry::Pio shows no recovery in the lumen and only minor intracellular recovery compared to *wt* (Movie S2). Dumpy:eYFP showed no recovery after bleaching, similar to *wt* embryos. The scale bar represents 10 µm.

#### Figure 1

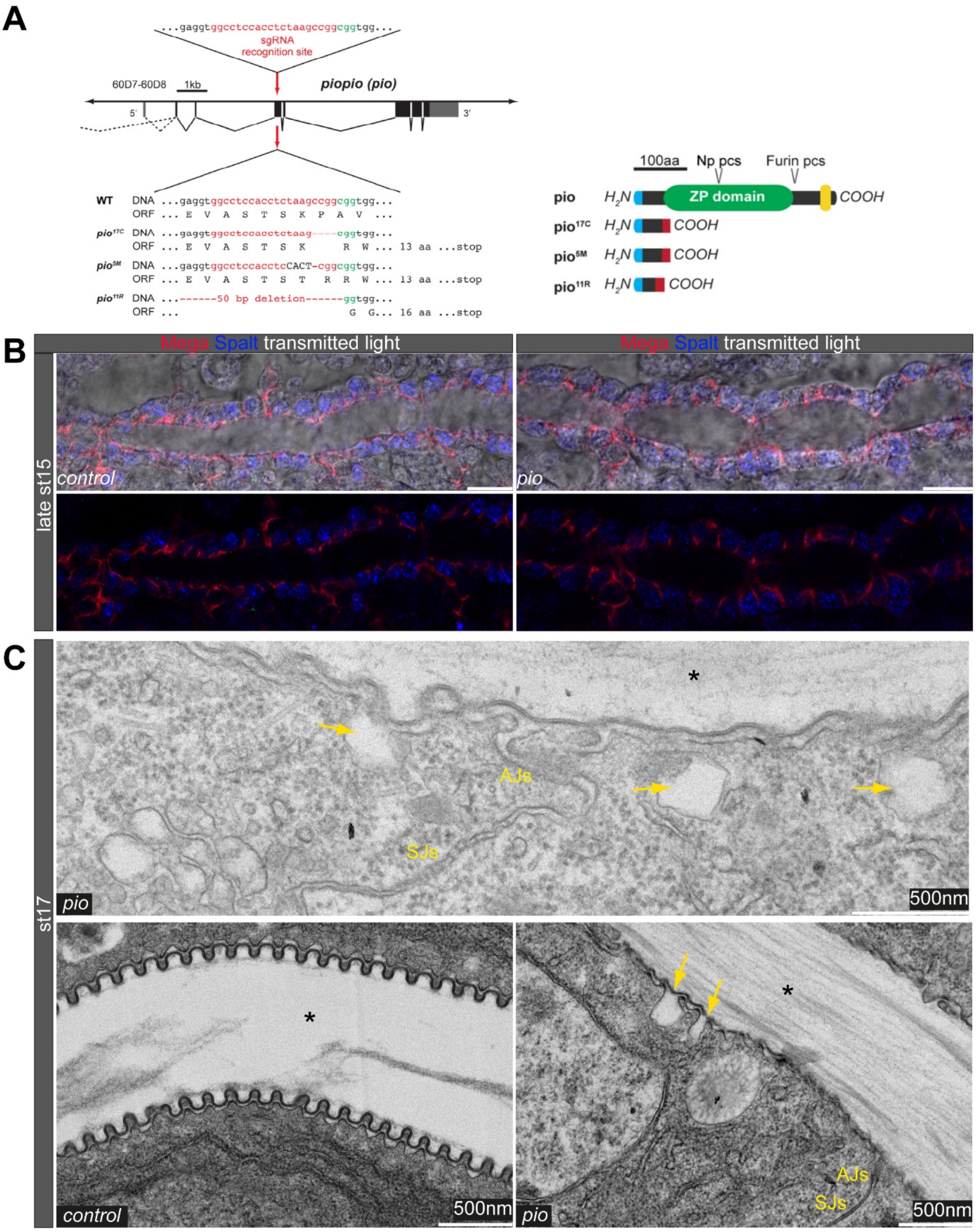


**Generation and ultrastructure analysis of *pio* mutant embryos**

**(A)** Generation of three independent *pio* mutant alleles which lack all relevant protein domains. Schema of CRISPR/Cas9-mediated mutagenesis to generate frame shift mutations in the 5' region of the pio ORF. Map of genomic *pio* region and of single guide RNA (sgRNA) target site (red letters) with PAM (green letters), ORF (black boxes) and UTRs (grey boxes). Alleles of *pio* with generated indels that cause ORF frame shifts in $pio^{17c}$, $pio^{5M}$ and $pio^{11R}$ mutant alleles. Frame shifts result in the expression of truncated Pio proteins. Signal peptide (blue), ZP domain (green), transmembrane domain (yellow), frame shifted amino acid sequence (red), and protease cleavage sites (pcs) are indicated.

SMART analysis of Pio (PB_FBpp0072219) supports previous findings by Jaźwińska, Ribeiro, and Affolter (2003) N-terminal ZP module (66-351aa) and C-terminal transmembrane domain (TM, 410-432aa) are indicated. The anti-Pio antibody (kindly provided by the Affolter lab) recognizes a polypeptide stretch between 186 and 200aa within (arrow) the ZP module. Thus, anti-Pio antibody detects Pio after Furin (predicted site at 355aa) and Np processing.

**(B)** Confocal images of *wt* and *pio* mutant stage 15 embryos. The lateral membrane of dorsal trunks is marked with Megatrachea (Mega; red, arrows), and transmission light visualizes the tracheal cells and lumen. In addition, the transcription factor Spalt (blue) is detectable in nuclei of *wt* and *pio* mutant tracheal cells of the dorsal trunk. Lower panels show Mega and Spalt expression in tracheal cells. Scale bars indicate 10µm.

**(C)** TEM analysis of late-stage 17 *wt* embryos. The control embryo is shown in the lower panel left-hand image. The *pio* mutant embryos (upper and lower right-hand images) show normal AJs and SJs formation. However, *pio* mutant embryos contained bulges of the apical cell membrane (yellow arrow), and the pattern of taenidial folds was disorganized. Note that differences of taenidial fold disorganization phenotypes (compare with Fig. 1D) can reflect the variations of individuals of stage 17 embryos. The luminal extracellular matrix material within the tube lumen is indicated (*). Magnification is indicated in the images with scale bars.

**Figure 2**

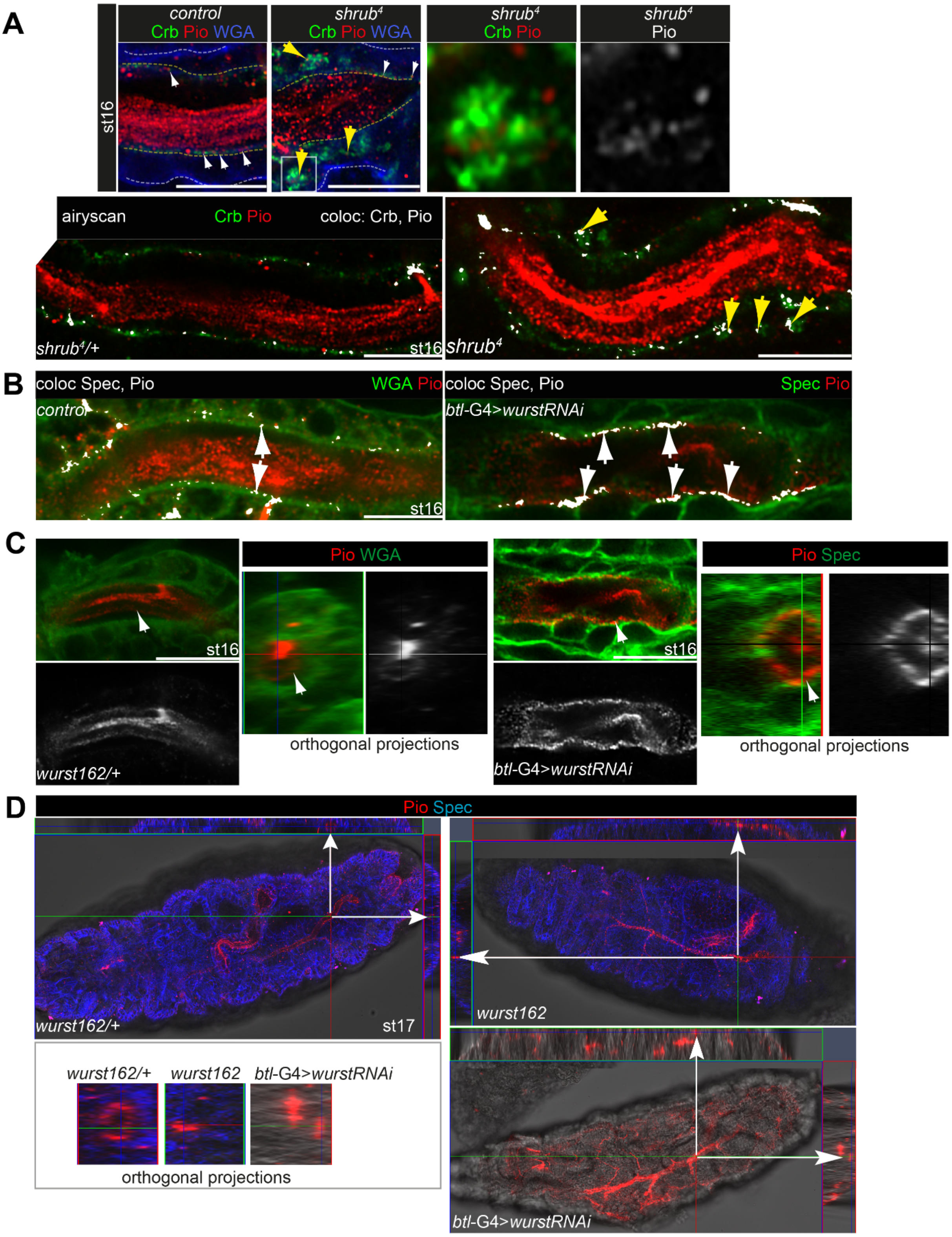


**Tube luminal matrix and Pio turnover in tracheal cells.**

**(A)** Confocal images of Pio and Crb in control and *shrub* mutant embryos and analyzed with the ZEN co-localization tool (lower panel, co-localization is indicated with white puncta). Pio is shown in red, Crb in green and overlap of both proteins in white in st16 embryos. Pio puncta overlap with Crb at the apical cell membrane (white arrows) in control and shrub mutant embryos and additionally in *shrub* mutants intracellularly in Crb marked swollen endosomes (yellow arrows).

**(B)** Confocal images of Pio, wheat germ agglutinin (WGA) and α-Spectrin in control and btl-Gal4 driven UAS-*wurst* RNAi knockdown st16 embryos, analyzed with the ZEN co-localization tool (co-localization is indicated with white puncta). Pio is shown in red, WGA and α-Spectrin in green and overlap in white. Control embryos show Pio puncta at the apical cell surface (white arrows). *wurst* knockdown embryos show accumulation of Pio puncta at the apical cell membrane (white arrows).

**(C)** Confocal images and orthogonal projections of the tube lumen after "express" deconvolution with SVI Huygens pro (*wurst*$^{162}$/+) and Airyscan mode (*wurst* RNAi). Pio is shown in red, WGA in green and α-Spectrin in green. White arrows point to Pio accumulation at the apical cell surface upon *btl*-Gal4>UAS-wurst RNAi knockdown.

**(D)** Confocal images and corresponding (white arrows) orthogonal projections of whole mount stage 17 *wurst* mutant, heterozygous control siblings and tracheal specific *btl-Gal4>UAS-wurst* RNAi knockdown embryos. Pio is shown in red, the cell membrane marker α-Spectrin in blue. While control embryos clear Pio from their tracheal tube lumen, *wurst* mutant and *wurst* knockdown embryos failed to clear luminal Pio. Magnifications of the orthogonal projections are shown left in the lower panel. Late st17 *wurst*$^{162}$ heterozygous control embryos show Pio at the apical cell surface but not within the tracheal tube lumen. Comparable *wurst*$^{162}$ mutant and wurst RNAi knockdown embryos show Pio staining within the tube lumen. Scale bars indicate 10µm.

#### Figure 3

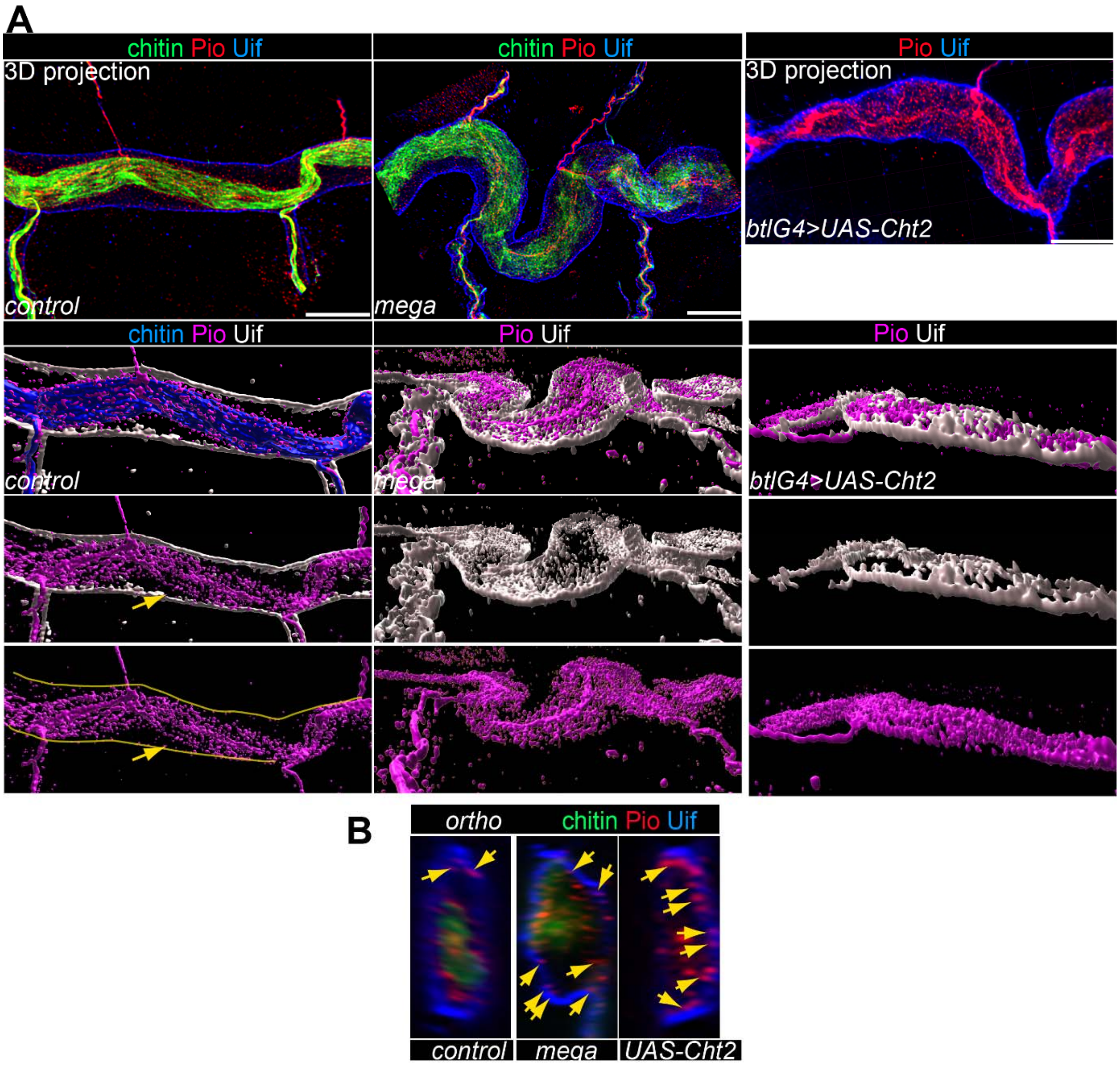


**Pio localization in *mega* mutant and Cht2 overexpressing embryos**

**(A,B)** 3D images (upper panel in A) of dorsal trunks and orthogonal projections (B) of airyscan Z-stacks show Pio (red), Uif (blue), and chitin (chitin-binding probe; green). Lower panels in A show 3D Visualization of Imaris surfaces of Voxel data resulting from airyscan Z-stacks, Pio (magenta), Uif (grey), and chitin-binding probe (blue). Control embryos show Pio staining accumulation in the lumen and some puncta (yellow arrows) at and near the apical cell membrane (yellow line in A). Mega mutant and Cht2 overexpressing embryos show luminal Pio but also Pio puncta enrichment at and near the apical cell membrane. Arrows in B point to Pio puncta at and near the apical cell membrane marked with Uif. Scale bars indicate 10μm.

#### Figure 4:

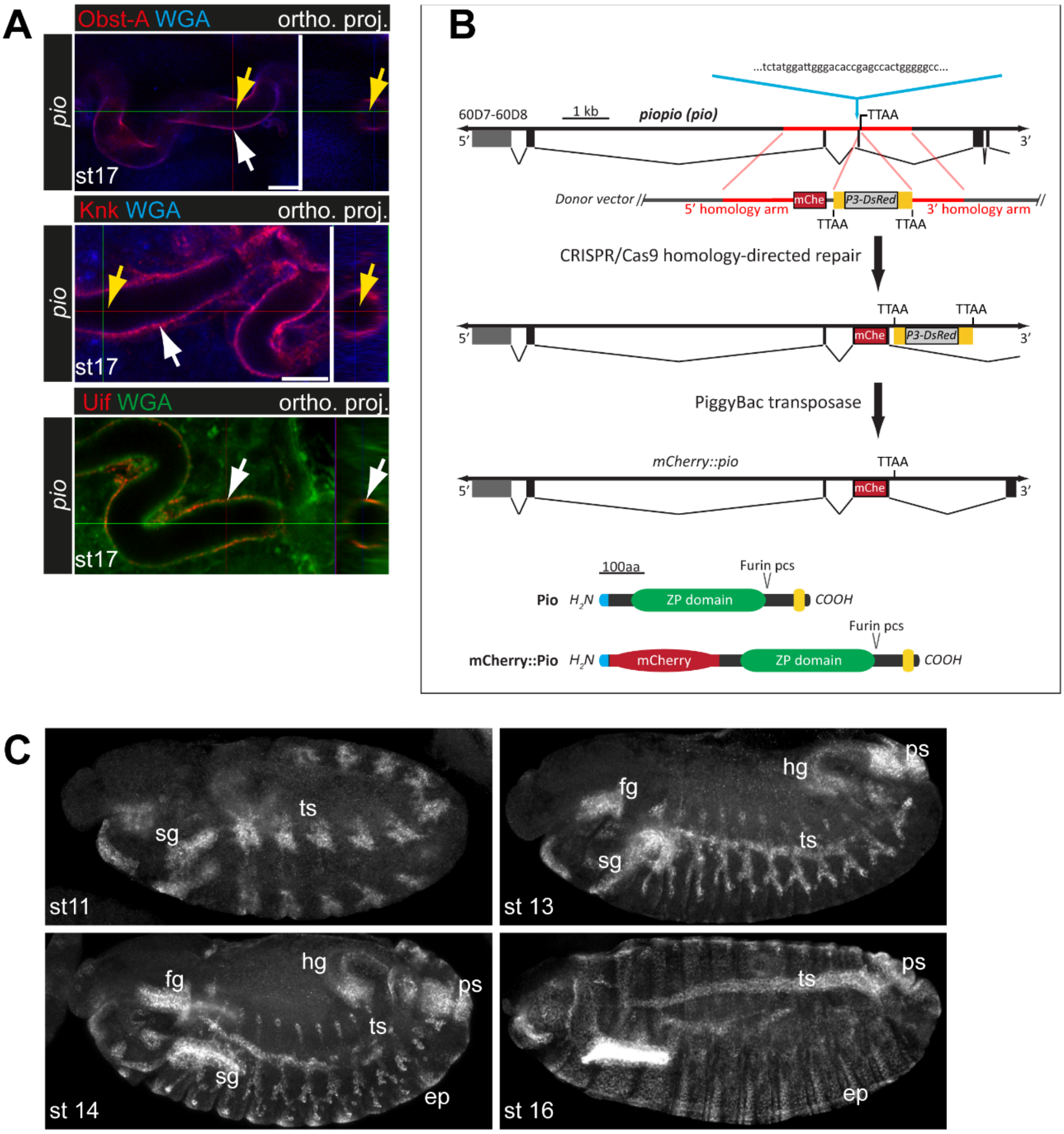


**CRISPR-Cas9 mediated *pio* mutagenesis and caused phenotypes and mCherry:Pio generation and expression**

**(A)** The stage 17 *pio* loss of function embryos are able to clear chitin-matrix proteins from tracheal tube lumina, as indicated by Obst-A (red), Knk (red) and WGA (blue) stainings. Shown are confocal overview and higher magnification images and the corresponding orthogonal projections of the tube lumen (right hand). White arrows point to the apical cell membrane and yellow arrows to the tube lumen area from which orthogonal projections were generated. Scale bars represent 10µm.

**(B)** Schema of the 5’ region of the pio genomic locus together with the donor vector containing two homology arms (red) and mCherry encoding sequences followed by a 3xP3-DsRed marker gene flanked by PiggyBac transposon ends (yellow). The sgRNA recognition site (magenta), PAM (green), translated DNA (black boxes) and UTRs (grey boxes) are indicated before and after CRISPR/Cas9-mediated homology-directed repair. Signal peptides (blue), mCherry (red), ZP domains (green), Furin cleavage sites (pcs) and transmembrane domains are indicated below in Pio and mCherry::Pio proteins.

**(C)** Confocal maximum intensity projections of anti-mCherry immunostainings of whole mount embryos. mCherry::Pio shows a Pio-characteristic ectodermal expression pattern. ep: epidermis, fg: foregut, hg: hindgut, ps: posterior spiracle, ts: tracheal system, sg: salivary glands.

**Figure 5:**

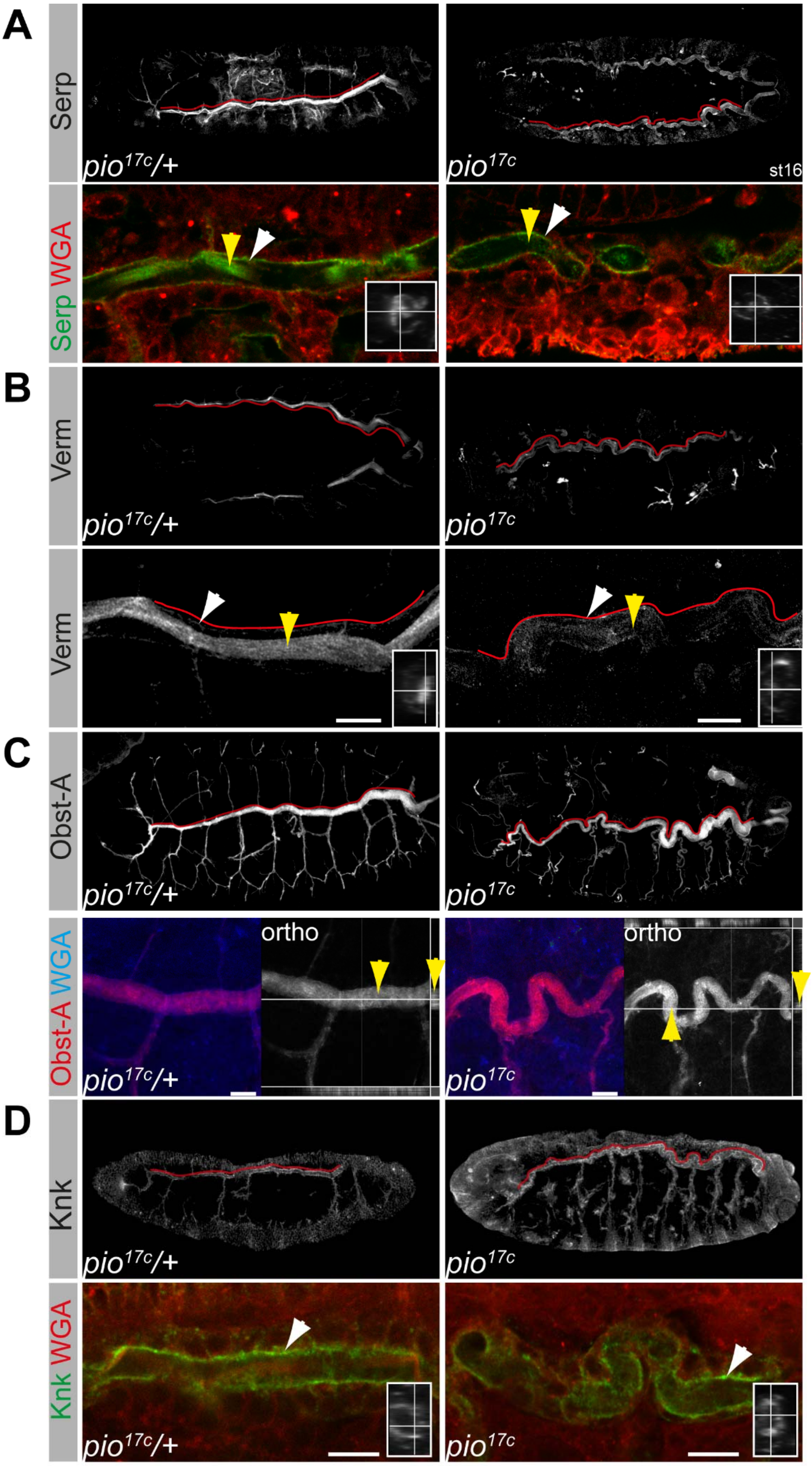
A
Serp
pio17c/+
pio17c
st16
Serp WGA
B
Verm
pio17c/+
pio17c
Verm
C
Obst-A
pio17c/+
pio17c
Obst-A WGA
ortho
pio17c/+
ortho
pio17c
D
Knk
pio17c/+
pio17c
Knk WGA
pio17c/+
pio17c

**Localization of Chitin-matrix proteins in *pio* mutant embryos**

Confocal images and maximum intensity projections (3D) of *pio* loss of function (right) mutant stage 16 embryos and heterozygous control siblings (left). Yellow arrows point to luminal staining; white arrows to staining at the apical surface. Dorsal trunk is indicated by red line. Scale bars represent 10µm

**(A, B)** Serp is shown in green, WGA in red, and single channels in grey. Overview and magnifications of the trachea (lower rows) reveal normal Serp (green) localization in *pio* mutants. Overview and magnifications of Verm reveal normal Verm (grey) localization within the chitin-matrix of pio mutant embryos. Framed inlays are orthogonal projections showing Serp or Verm within the tracheal lumen. Note the reduced luminal Serp and Verm staining (yellow arrows) in *pio* mutant embryos when compared to the surface staining (white arrow) and control embryos.

**(C, D)** Z-stack projections of tracheal Obst-A and Knk stainings. Overview (upper) and magnifications (lower rows) reveal normal subcellular Obst-A and Knk localization within the chitin matrix of *pio* mutant embryos. Framed inlays are orthogonal projections showing Knk within the tracheal lumen when compared with the control embryos. Ortho (in C); orthogonal projection of X and Y axis showing Obst-A within the lumen. Scale bars indicate 10µm.

#### Figure 6

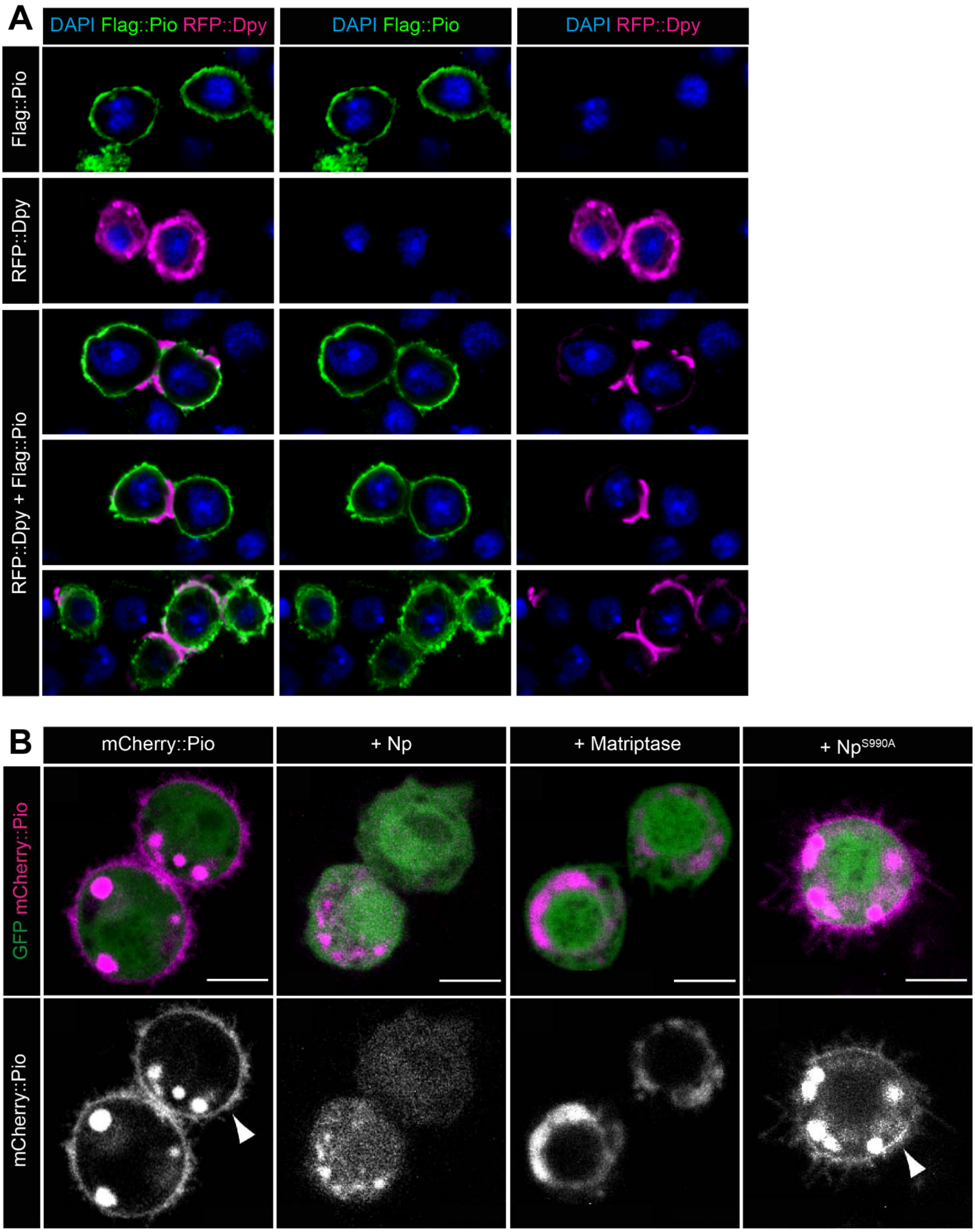


**Pio is involved in Dpy secretion and Notopleural and matriptase activity controls Pio localization.**

**(A)** Confocal images of single or co-expression of FLAG-tagged Pio, and RFP-tagged Dpy C-terminal region (ZPD domain, transmembrane domain, cytoplasmic region) in the *Drosophila* S2R+ (Schneider) cells. Single FLAG-tagged Pio expression reveals Flag::Pio staining at the cell surface. Singly RFP-tagged Dpy expression revealed predominant intracellular staining of RFP::Dpy. Co-expression of FLAG-tagged Pio and RFP-tagged Dpy revealed predominant RFP::Dpy staining at the cell surface colocalizing with Flag::Pio. See Figure 4 for a schematic representation of the expressed recombinant proteins. Here, the RFP::Dpy$^{ZP}$ construct was used but the same results were also observable with the RFP::Dpy$^{CT}$ construct (not shown)

**(B)** Live cell confocal images show UAS/Gal4 mediated GFP (green) and mCherry::Pio (magenta) expression in *Drosophila* Kc167 cells. Control cells (left panel) reveal mCherry::Pio signals predominantly at the cell surface (arrowhead). Upon co-expression of Np or Matriptase, cell surface mCherry::Pio signal was lost (middle panels). The co-expression of the catalytical inactive Np$^{S990A}$ mutant did not affect extracellular mCherry::Pio signals (arrowhead, right panel).

169 **Figure 7**

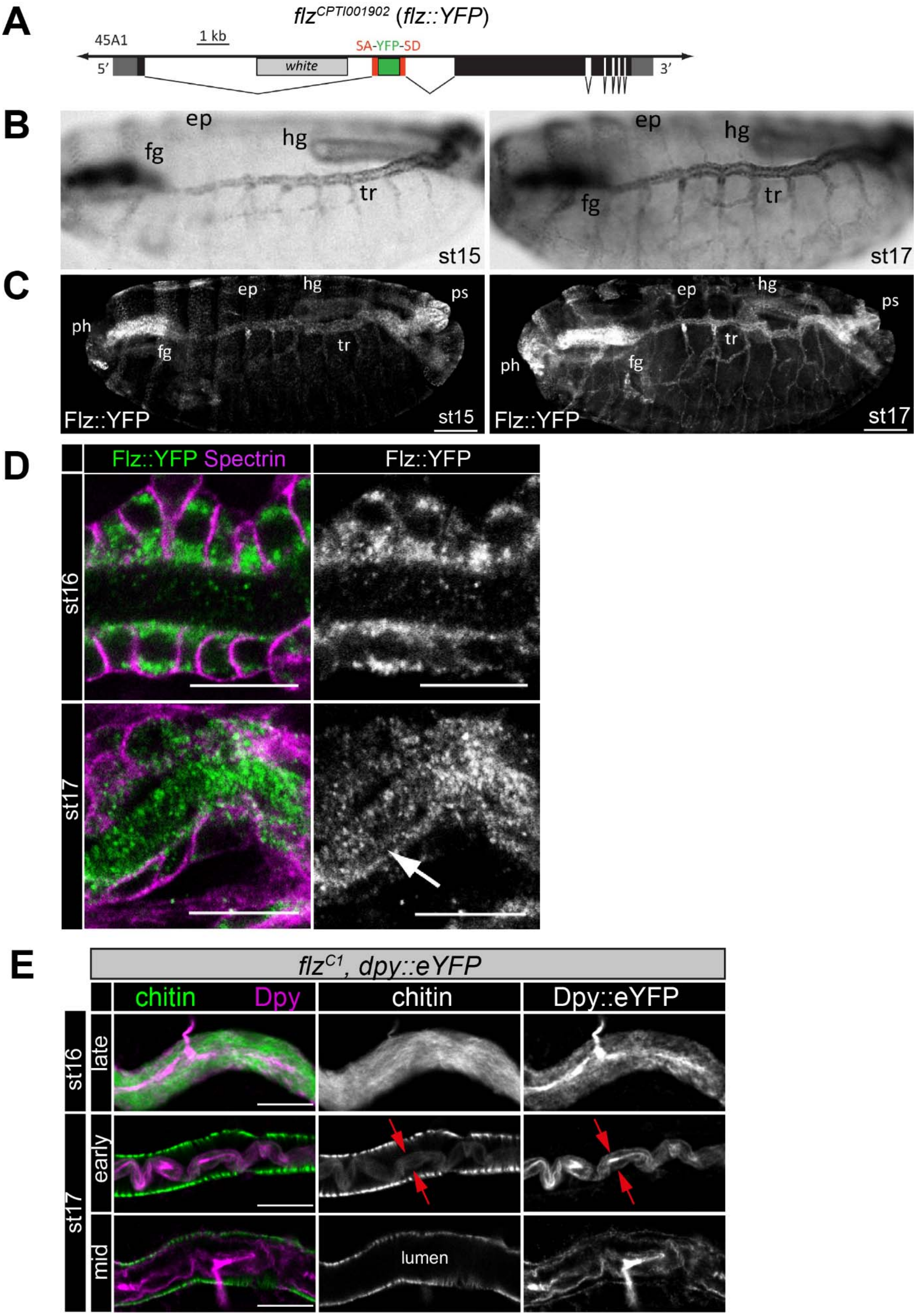


170

171 **The tracheal expression and localization of the serine protease Filzig (Flz).**

**(A)** The *flz* allele includes YFP coding sequence (green box) flanked by splice acceptor (SA) and donor (SD) sites.

**(B)** Bright field microscopy of stainings of *in situ* RNA hybridization of DIG-labeled *flz* antisense probe. The stage 16 and 17 embryos show tracheal *flz* RNA expression in ectodermal epithelia (ep, epidermis; fg, foregut; hg, hindgut; tr, trachea).

**(C,D)** Embryonic Flz::YFP expression (C) resembles *flz* RNA pattern (B). Confocal images of anti GFP (green) and anti $\alpha$-Spectrin (magenta) antibody stainings reveal predominant Flz::YFP localization within the tracheal lumen (arrow) of stage 17 embryos but not at stage 16. Scale bars represent 50 µm (C) or 10 µm (D).

**(E)** Confocal images of dorsal trunks of *flz* mutant stage 16 (upper), early stage 17 (middle) and mid stage 17 (lower panel) embryos with endogenous expression of Dpy::eYFP stained with anti GFP antibody (magenta) and cbp (chitin; green). Single channels are indicated in grey. The *flz* mutant embryos reveal normal Dpy release and extracellular localization but no clearance of luminal Dpy as in *wt* embryos (see Movie S1). However, the luminal Dpy and chitin cable condenses (red arrows) during chitin degradation as in *wt* (compare with Fig 6A). Scale bars represent 10 µm**.**

#### Figure 8

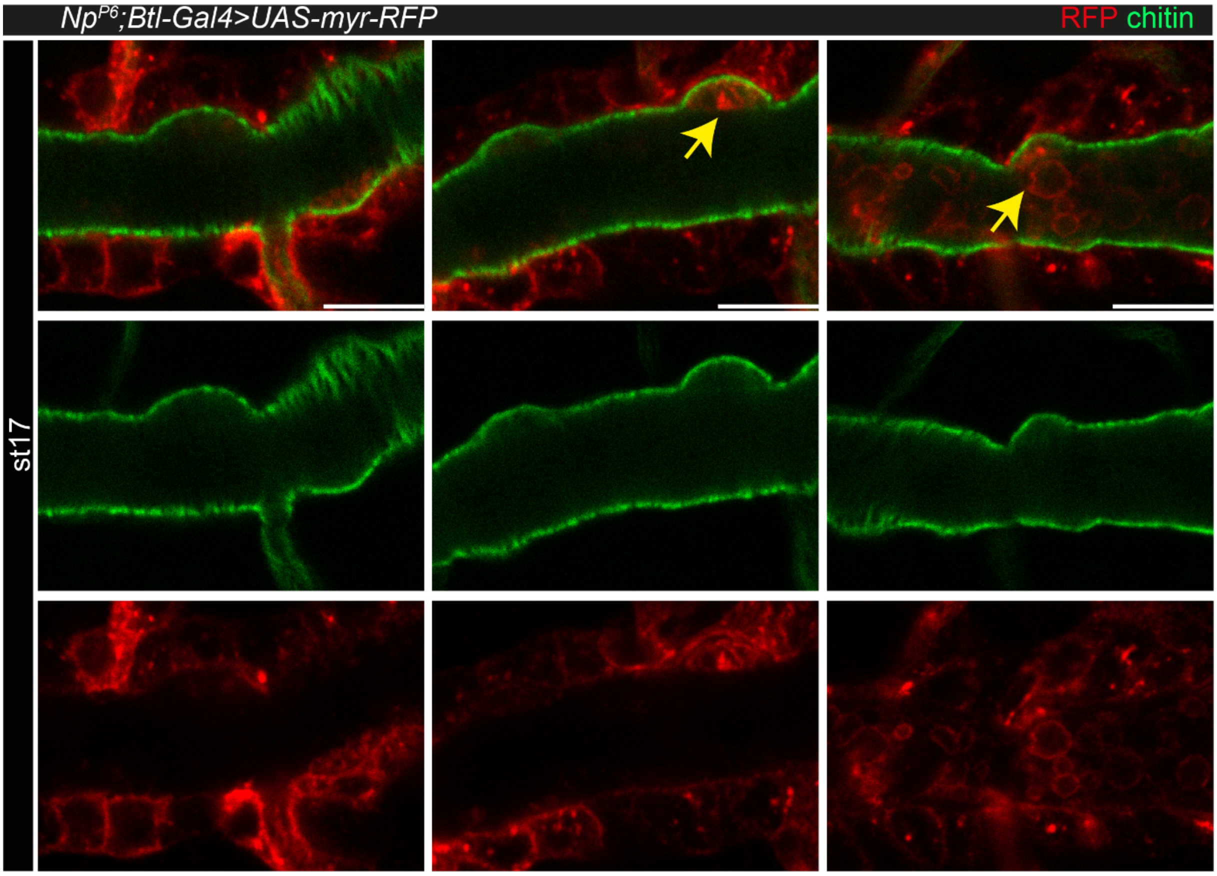


**Tracheal expression of myr-RFP in *Np* mutant embryos**

Confocal images of *btl-Gal4* driven expression of *UAS-myr-RFP* in *Np* mutant embryos. Antibody stainings show RFP (red) and chitin (cbp; green) in mid to late (left to right) stage 17 embryos. Arrows point to unusual luminal RFP staining, detected first at the aECM bulge and subsequently in the tube lumen. Scale bars indicate 10µm.

#### Figure 9

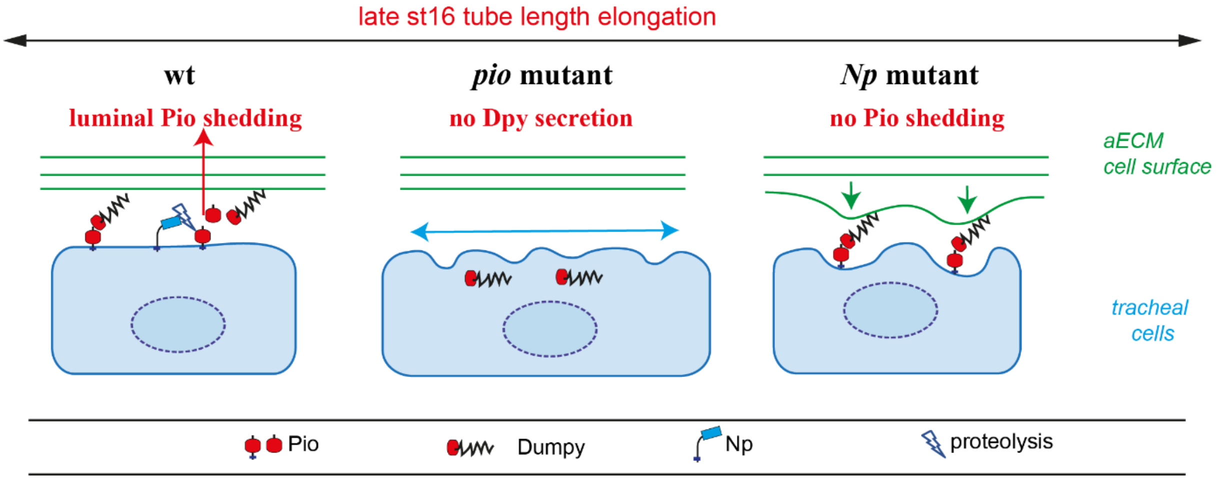


**Model of apical Pio Dpy matrix, Pio ectodomain shedding, and Dpy secretion**

The model shows a Pio-mediated Dpy secretion, the apical ZP domain protein matrix formation, and subsequent Pio ectodomain shedding by Np in wt tracheal cells. In *pio* null mutant embryos, Dpy is not secreted. The loss of the apical ZP proteins led to membrane matrix disconnection. This results in unstable membrane structures with numerous unusual gaps between membrane and matrix, excess apical cell shape expansion (indicated by the blue double arrow) in the axial direction, and dorsal trunk overexpansion. In contrast, apical ZP domain protein matrix forms in Np mutant embryos, but apical Pio ectodomain shedding is prevented. This led to membrane bulging and rupture (indicated by green arrows) of the apical ECM at bulges and slightly shortened dorsal trunks.

### Materials and Methods

#### List of used antibodies, fly strains, oligos and software

| Antibodies and reagents | Dilution | Source | Identifier |
|---|---|---|---|
| anti-Crb, mouse | 1:10 | DSHB | Cq4 |
| anti-Flag, mouse | 1:500 | Merck | F3165 |
| anti-Gasp/Obst-C, mouse | 1:5 | DSHB | 2A12 |
| anti-GFP, chicken | 1:1000 | Abcam | ab13970 |
| anti-GFP, rabbit | IF: 1:500<br>WB: 1:10000<br>CIF: 1:1000 | Synaptic Systems | 132003 |
| anti-Knk, rabbit | 1:50 | Moussian | (Moussian et al., 2006) |
| anti-Mega, mouse | 1:50 | Schuh | (Jaspers et al., 2012) |
| anti-mCherry, rabbit | IF: 1:500<br>WB: 1:10000 | Rockland | 600-401-P16 |

| | | | |
|---|---|---|---|
| anti-Obst-A, rabbit | 1:50 | Behr | (Petkau et al., 2012) |
| anti-Pio, rabbit | 1:100 | Affolter | (Jaźwińska et al., 2003) |
| anti-Serp, rabbit | 1:50 | Luschnig | (Luschnig et al., 2006) |
| anti-Spalt, rabbit | 1:25 | Schuh | (Kuhnlein et al., 1994) |
| anti-α-Spectrin, mouse | 1:10 | DSHB | 3A9 |
| anti-Strep-HRP, mouse | 1:10000 | IBA | 1-1509-001 |
| anti-β-Tubulin, mouse | 1:100 | DSHB | E7 |
| ani-Uif, guinea pig | 1:100 | Ward | (Zhang and Ward, 4th., 2009) |
| anti-Verm, rabbit | 1:50 | Luschnig | (Luschnig et al., 2006) |
| anti-goat Alexa488, donkey | 1:400 | Dianova | 705-545-003 |
| anti-guinea pig Cy3, donkey | 1:400 | Dianova | 706-165-148 |
| anti-mouse Alexa647, donkey | 1:400 | Dianova | 715-605-020 |
| anti-mouse Alexa647, donkey | 1:400 | Dianova | 715-605-020 |
| anti-rabbit Cy3, donkey | 1:400 | Dianova | 711-167-003 |
| anti-rabbit-AlexaFluor488 | 1:500 | Thermo Fisher Scientific | A-11034 |
| anti-rabbit-AlexaFluor568 | 1:500 | Thermo Fisher Scientific | A-21069 |
| anti-mouse-HRP | WB: 1:10000 | Thermo Fisher Scientific | G-21040 |
| anti-rabbit-HRP | 1:10000 | Thermo Fisher Scientific | G-21234 |
| WGA, Alexa Flour 633 | 1:100 | Invitrogen | W21404 |
| Cbp, Alexa488 | 1:200 | New England Biolabs | |
| **fly strains** | | | |
| **Strain** | | **Source** | **Identifier** |
| btl-Gal4 | | BDSC | |
| $crb^{2}$ ($crb^{11A22}$) | | BDSC | Stock ID 3448 |
| *$mega^{G0012}$/FM7, act-GFP* | | Schuh; Schäfer | (Behr et al., 2003) |
| *$shrub^{4}$*/Cyo | | S. Hayashi | (Dong et al., 2014) |
| *$w^{1118}$* | | BDSC | |
| *$w^{*}$; mCherry::pio/CyO, dfd-eYFP* | | Drees | This manuscript |
| *$w^{*}$; $flz^{C1}$* | | Drees | This manuscript |
| *$w^{*}$; $flz^{C1}$, mCherry::pio/CyO, dfd-eYFP* | | Drees | This manuscript |
| *$w^{1118}$; PBac{681.P.FSVS-1}$flz^{CPTI001902}$* | | Kyoto stock center | Stock ID 115246 |
| $w^{*}$; $pio^{5M}$/CyO, dfd-eYFP | | Drees | This manuscript |
| $w^{*}$; $pio^{17C}$/CyO, dfd-eYFP | | Dress | This manuscript |
| $w^{*}$; $Np^{P6}$/CyO, dfd-eYFP | | Drees | (Drees et al., 2019) |
| $w^{*}$; $Np^{P6}$, P{Gal4-btl}/CyO, dfd-eYFP | | Drees | (Drees et al., 2019) |
| $w^{*}$; $Np^{C2}$, P{UAS-$Np^{S990A}$}/CyO, dfd-eYFP | | Drees | (Drees et al., 2019) |
| $w^{*}$; $Np^{C2}$/CyO, dfd-eYFP ; P{UASp-GFPS65C-alphaTub84B}3/TM3, $Sb^{1}$ | | Drees | (Drees et al., 2019) |
| $w^{*}$; $Np^{P6}$, mCherry::pio/CyO, dfd-eYFP | | Drees | (Drees et al., 2019) |
| *$w^{*}$ ; P{UAS-$Np^{S990A}$}/P{UAS-$Np^{S990A}$}* | | Drees | (Drees et al., 2019) |
| *$w^{*}$ ; P{UAS-Np}/P{UAS-Np}* | | Drees | (Drees et al., 2019) |
| *$wurst^{162}$*/FM7-actin-GFP | | Behr | (Behr et al., 2007) |
| *UAS-Cht2* | | Uv | (Tonning et al., 2005) |
| *w[1118]; P{w[+mC]=UAS-myr-mRFP}1* | | BDSC | Stock ID 7118 |

| | | |
|---|---|---|
| *UAS-wurst-RNAi* | VDRC | (Stümpges and Behr, 2011) |
| **Oligonucleotides** | | |
| **Identifier** | **Sequence** | **Source** |
| pio-sgRNA-sense | CTTCGATTGGGACACCGAGCCACT | Eurofins Genomics |
| pio-sgRNA-antisense | AAACAGTGGCTCGGTGTCCCAATC | Eurofins Genomics |
| flz-sgRNA-sense | CTTCGTGGGTTACGCCGG CCTCAA | Eurofins Genomics |
| flz-sgRNA-antisense | AAACTTGAGGCCGGCGTA ACCCAC | Eurofins Genomics |
| UAS-mCherry::pio-for | GAATTCATGAAGACAGGCACTCGAATGGACGCTTT CCACACGGCGCTGCACTTAATCACAATCGCAGCTC TGACGACG | Eurofins Genomics |
| UAS-mCherry::pio-rev | CTCGAGGCCGCCTTTGTAAAGCTCATCC | Eurofins Genomics |
| Pio-5'-HA1-for | ACTAGTCCGAATTCGCAGG TGATTATCGCCTCTCGGCC ATCAG | Eurofins Genomics |
| Pio-5'-HA1-rev | AAGCTTCTTTAATTAAAGG GGAAATTTCG | Eurofins Genomics |
| Pio-5'-HA2-for | ACTAGTGGCAAGCTTACTG GCGATGGATTAGGCC | Eurofins Genomics |
| Pio-5'-HA2-rev | CACCTGCGATCTTAATCTT GCCAGCGTCTGTC | Eurofins Genomics |
| Pio-3'-HA-for | TTAAGGAAGAGCACACAG TTGGGCGCTTTGTTAGTCG | Eurofins Genomics |
| Pio-3'-HA-rev | CGGGGAAGAGCGACGAGA TTGCGCCGGAAAATAAG | Eurofins Genomics |
| UAS-pio-ORF-for | CTCGAGCCAACGGCAATGAAAGATGCCC | Eurofins Genomics |
| UAS-pio-ORF-rev | TCTAGATTAGCTGCTGTGCGAGAAG | Eurofins Genomics |
| Dpy-ZP-for | GCTTTACAAAGGTTACACGGGTAATCCG | Eurofins Genomics |
| Dpy-CT-for | GCTTTACAAAGGTGGAAATGCCAGGATTG | Eurofins Genomics |
| Dpy-CTZP-rev | GTGGAGCCGGCCACCATTTATGGAGGTTTC | Eurofins Genomics |
| Dpy-ZP-for | GGCCACCATTTATGGAGGTTTC | Eurofins Genomics |
| Dpy-ZP-rev | GGTTCCTTCACAAAGATCCTTTAGGATATGTAATCC GGCG | Eurofins Genomics |
| Strep::TGFBR3::GFP1-for | CTGAATAGGGAATTGGGAATTCATGACTTCCCATTA TG | Eurofins Genomics |
| Strep::TGFBR3::GFP1-rev | CACCGCTGCCACCTCCTGATCCGCCACCCTTTTCA AACTGCGGATGACTCCATGCACTTTGCACCTCTTCT GGCTCTC | Eurofins Genomics |
| Strep::TGFBR3::GFP2-for | ATCAGGAGGTGGCAGCGGTGGAAGTGCATGGAGC CATCCCCAATTCGAGAAGGGGAGCGTGGATATTGC CCTG | Eurofins Genomics |
| Strep::TGFBR3::GFP2-rev | TCACCATACCGCCGCTAGCGGCCGTGCTGCTGCT G | Eurofins Genomics |
| **cloning** | | |
| **Identifier** | **Purpose** | **Source** |
| pJet1.2 | Cloning of amplicons | Thermo Fisher Scientific |
| pUAST | GAL4/UAS-mediated overexpression | (Brand and Perrimon, 1993) |
| pBFv-U6.2 | Expression of single sgRNA | (Kondo and Ueda, 2013) |
| pBFv-U6.2B | Expression of two sgRNAs | (Kondo and Ueda, 2013) |

| | | |
|---|---|---|
| pHD-ScarlessDsRed | Scarless genome editing via HDR | DGRC |
| actin5C-Gal4 | Expression of Gal4 in cultured cells | (Usui et al., 1999) |
| **Software and algorithms** | | |
| Clustal omega algorithm | | https://www.ebi.ac.uk/Tools/msa/clustalo/ |
| DNASTAR software suite | Lasergene Software | |
| Flybase | www.flybase.org | BLASTP algorithm |
| Huygens professional | SVI | 20.10 |
| Illustrator | Adobe | CS6 |
| Imaris 9.7.2 | Oxford Instruments | https://imaris.oxinst.com/ |
| NetOGlyc | | https://services.healthtech.dtu.dk/ |
| Office 365 (Word, Excel) | Microsoft | https://www.microsoft.com |
| ProP | | https://services.healthtech.dtu.dk/ |
| SignalP | | https://services.healthtech.dtu.dk |
| Photoshop CS6 | Adobe | https://www.adobe.com |
| SMART | EMBL | |
| Serial Cloner | Serial basics | 2.6.1 |
| TMHMM 2.0 algorithm | | https://services.healthtech.dtu.dk/ |
| ZEN 2.3 | Zeiss | 2.3, black |
